# Synergistic reinforcement learning by cooperation of the cerebellum and basal ganglia

**DOI:** 10.1101/2024.07.12.603330

**Authors:** Tatsumi Yoshida, Hikaru Sugino, Hinako Yamamoto, Sho Tanno, Mikihide Tamura, Jun Igarashi, Yoshikazu Isomura, Riichiro Hira

**Affiliations:** Department of Physiology and Cell Biology, Graduate School of Medical and Dental Sciences, Institute of Science Tokyo, Tokyo, Japan; High Performance Artificial Intelligent System Research Team, Center for Computational Science, RIKEN, Saitama, Japan

## Abstract

The cerebral cortex, cerebellum and basal ganglia are essential for flexible learning in mammals. Although traditionally thought to operate under different learning rules, recent evidence suggests that both the basal ganglia and the cerebellum may employ reinforcement learning mechanisms. This raises the question of how these structures coordinate when a common reward prediction error mechanism is active. To address this issue, we first examined output signals from the basal ganglia and cerebellum following the activity of the cerebral cortex. We recorded single-neuron activity from the output regions of the cerebellum and basal ganglia - the cerebellar nuclei (CN) and substantia nigra pars reticulata (SNr) - in both male and female ChR2 transgenic rats. Neurons in the CN and SNr exhibited distinct temporal response patterns; notably, the fast excitatory response in the CN, driven by mossy fiber input, was synchronized with the inhibitory response in the SNr, mediated via the direct pathway. Using these experimental findings together with connectome data, we developed both a semi-realistic spiking network model and a reservoir-based reinforcement learning model. In the latter model, successful learning depended on synaptic plasticity in both the cerebellum and basal ganglia with a temporal precision on the order of 10 ms. Furthermore, cortical β-oscillations enhanced learning and optimal reinforcement learning occurred when the output of cerebellar and basal ganglia signal phase-locked at the frequency of cortical oscillation. Taken together, our results suggest that the coordinated output of the cerebellum and basal ganglia, driven by tightly tuned cortical input, underlies brain-wide synergistic reinforcement learning.

**Significance Statement:** The cerebral cortex, cerebellum, and basal ganglia support learning. Recent research suggests that both the basal ganglia and cerebellum use a similar learning process called reinforcement learning, which involves predicting rewards. To understand how these brain regions work together, we recorded brain activity in rats while photo-stimulating the cerebral cortex. We found that two types of responses in the cerebellum and basal ganglia were synchronized, which might help activate the cerebral cortex. A computer model showed that precise timing of signals from both the cerebellum and basal ganglia is important for learning. This timing was important only when the cerebral cortex worked in a specific frequency range. Our findings suggest that coordinated brain activity enhances learning.

## Introduction

The brain is a complex system composed of many unique information-processing regions that operate in a coordinated manner to produce flexible and sophisticated behavior (Caligiore et al. 2019; Doya 2000; Caligiore et al. 2017; Lake et al. 2017; Penhune and Steele 2012; Shadmehr and Krakauer 2008). To understand the mechanism of cooperative information processing, it is necessary to elucidate how the multiple interregional loop circuits work together. The cerebral cortex outputs to numerous subcortical structures, including the cerebellum and basal ganglia. These regions differ in many evolutionary and functional aspects but are similar in the fact that they send signals back to the cerebral cortex via the thalamus (Grillner 2021; Kebschull et al. 2020; Allen and Tsukahara 1974; Alexander and Crutcher 1990; Schell and Strick 1984). These multi-synaptic pathways, each with its own specific microcircuits and integration strategies, substantially affect ongoing activity of the cerebral cortex and entire brain.

Cerebellar nuclei (CN) are the output nuclei of the cerebellum and have direct thalamic projections, receiving inhibitory inputs from Purkinje cells (PCs), which are activated by excitatory granule cells that receive excitatory mossy fiber (MF) inputs. MFs are activated directly from the cerebral cortex, and consequently many neurons in CN receive inhibitory inputs from the cerebral cortex. MFs also directly excite CN neurons (Sugihara, Wu, and Shinoda 1996; Shinoda et al. 1992). Thus, CN receive multi-phase excitatory and inhibitory input from the cerebral cortex. The substantia nigra pars reticulata (SNr), an output nucleus of basal ganglia, is composed of inhibitory neurons projecting to the thalamus. In the SNr, a triphasic response to cerebral cortex stimulation in the order of excitation, inhibition, and excitation has been characterized in detail (Nambu et al. 2000; Chiken, Shashidharan, and Nambu 2008). To understand how these responses with excitatory and inhibitory dynamics are reintegrated in the thalamus and cerebral cortex, the relationships between multiple cerebral cortex areas and the subdivisions within cerebellum and basal ganglia should be systematically examined under identical conditions.

Coordination of cerebro-cerebellar loop plays an important role in cognitive and motor tasks (Wagner et al. 2019; Proville et al. 2014; Chabrol, Blot, and Mrsic-Flogel 2019; Gao et al. 2018; Zhu et al. 2023). Coordination of cerebro-basal ganglia thalamocortical loop is also critical in normal behavior and in pathological conditions (Wang et al. 2021; Alexander, DeLong, and Strick 1986; Yoshizawa et al. 2023; Stein and Bar-Gad 2013; Yasoshima et al. 2005; Alexander and Crutcher 1990). Classically, the basal ganglia and cerebellum were thought to be responsible for reinforcement and supervised learning, respectively (Doya 1999). However, the cerebellum has recently been shown to be also capable of reinforcement learning, as complex spikes in PCs are associated with reward prediction errors (Sendhilnathan et al. 2020; Yamazaki and Lennon 2019; Kostadinov and Häusser 2022; Thoma et al. 2008; Heffley and Hull 2019; Hoang et al. 2023; Boven et al. 2023; Pemberton, Chadderton, and Costa 2022). Thus, plastic changes in parallel fiber input in PCs due to climbing fibers input and plastic changes in corticostriatal pathways due to dopamine release could both lead to reinforcement learning. As circuit changes generated by different mechanisms facilitate learning, they must be consistently reintegrated in the cerebral cortex via the thalamus for decision making and motor output.

The cerebellum and basal ganglia each exhibit unique oscillatory activities (Halje et al. 2019; Courtemanche, Robinson, and Aponte 2013). For example, cerebellar oscillations in the β band range (15–30 Hz) is crucial for regulating action timing in human (Spencer and Ivry 2021). γ oscillations (30–80 Hz) can be also generated by cerebellar cortical circuitry (Middleton et al. 2008). β oscillation is simultaneously enhanced in cerebral cortex and basal ganglia when determining motor output in rats (Leventhal et al. 2012). In motor cortex and basal ganglia, the β band oscillation is enhanced during reaching, grasping and other motor tasks (Khanna and Carmena 2015). The excessive β oscillation is observed in patients of Parkinson’s disease (Gambosi et al. 2024; Morén et al. 2019). Thus, both the cerebellum and basal ganglia exhibit oscillations in the β-γ range, and their proper timing and amplitude are essential for behavioral coordination. The close frequency band activity of the cerebellum and basal ganglia may contribute to the coordinated information processing of the entire brain (Tanaka, Kameda, and Okada 2024; Diedrichsen, Ivry, and Pressing 2003; Petter et al. 2016).

In this study, we investigated the propagation of neural activity patterns of the cerebellum and basal ganglia in response to cerebral cortex photoactivation using high-throughput optogenetic stimulation (Hira et al. 2013; Hira et al. 2015; Hira et al. 2009; Sugino et al., 2024) and neuronal activity recordings with the high-density Neuropixels probes (Jun et al. 2017). Although the temporal response patterns differed, CN excitatory signals and SNr disinhibitory signals were synchronously elevated. We modeled these circuits with a spiking neuronal network based on connectome data and a reservoir-based reinforcement learning model and analyzed these synergistic features. The results indicate that the synergy can facilitate brain-wide consistent reinforcement learning.

## Materials and Methods

### Animal preparation

All experiments were approved by the Animal Research Ethics Committee of the Institutional Animal Care and Use Committee of Tokyo Medical and Dental University (A2019-274) and were carried out in accordance with the Fundamental Guidelines for Proper Conduct of Animal Experiment and Related Activities in Academic Research Institutions (Ministry of Education, Culture, Sports, Science and Technology of Japan). We used 17 adult male and female rats (Thy1.2-ChR2-Venus transgenic rats, Long–Evans strain (Tomita et al. 2009; Saiki et al. 2018) at the age of 10-29 weeks for cerebellar recording and 10-14 weeks for SNr recording. These rats were kept under an inverted light schedule (lights off at 12 AM; lights on at 12 PM) in their home cages to adapt to experimental surroundings.

### Surgery

Primary surgery was performed to attach a custom-made head-plate (**Fig. 2C**) to the skull of rats under anesthesia by isoflurane gas (4.0–4.5% for induction and 2.0–2.5% for maintenance, Pfizer Japan, Tokyo, Japan) using an inhalation anesthesia apparatus (Univentor 400 anesthesia unit, Univentor, Zejtun, Malta). The body temperature was maintained at 37.0°C using an animal warmer (BWT-100, Bio Research Center, Aichi, Japan). The head of rats was fixed on a stereotaxic frame (SR-10R-HT, Narishige) with ear bars, and applied with lidocaine (Xylocaine Jelly, Aspen Japan, Tokyo, Japan) for local skin anesthesia and povidone-iodine disinfectant solution (10%, Kaneichi, Osaka, Japan) for disinfection around surgical incisions. The head-plate was then glued to the skull with stainless steel screws and dental resin cement (Super-Bond C & B, Sun Medical, Shiga, Japan; Unifast II, GC Corp., Tokyo, Japan). Reference and ground electrodes (PFA-coated silver wires, A-M systems, WA; 125-mm diameter) were implemented under the bone on the cerebellum. When we recorded the neurons in cerebellum, the reference electrode was placed over the anterior-most part of the cerebellum to avoid interference. Analgesics and antibiotics (meloxicam, 1 mg/kg sc, Boehringer Ingelheim, Tokyo, Japan; gentamicin ointment, 0.1% us. ext., MSD, Tokyo, Japan) were finally applied to remove pain and prevent infection.

More than three days later, secondary surgery was performed under the isoflurane anesthesia. For insertion of recording electrodes, we made a cranial window onto the cerebral cortex of left or right hemisphere or onto the cerebellum of the right hemisphere. The bone and dura mater were opened and removed by a dental drill (Tas-35LX, Shofu, Kyoto, Japan) and a dura picker (DP-T560-80, Bio Research Center, Aichi, Japan). The cortical surfaces were washed with PBS containing antibiotic (0.2% amikacin sulfate, Sawai, Osaka, Japan). We thinned the skull over the dorsal cerebral cortex and applied silicone oil to keep transparency for optogenetic stimulation (**Fig. 1F**). Thinned-skull area covered M1, M2, S1, and PPC. Because barrel cortex was not always completely covered with the thinned-skull window, we excluded the barrel cortex from S1 for the analysis when comparing the responsiveness. Analgesics (meloxicam, 1 mg/kg sc, Boehringer Ingelheim, Tokyo, Japan) was applied to remove pain.

**Figure 1.**
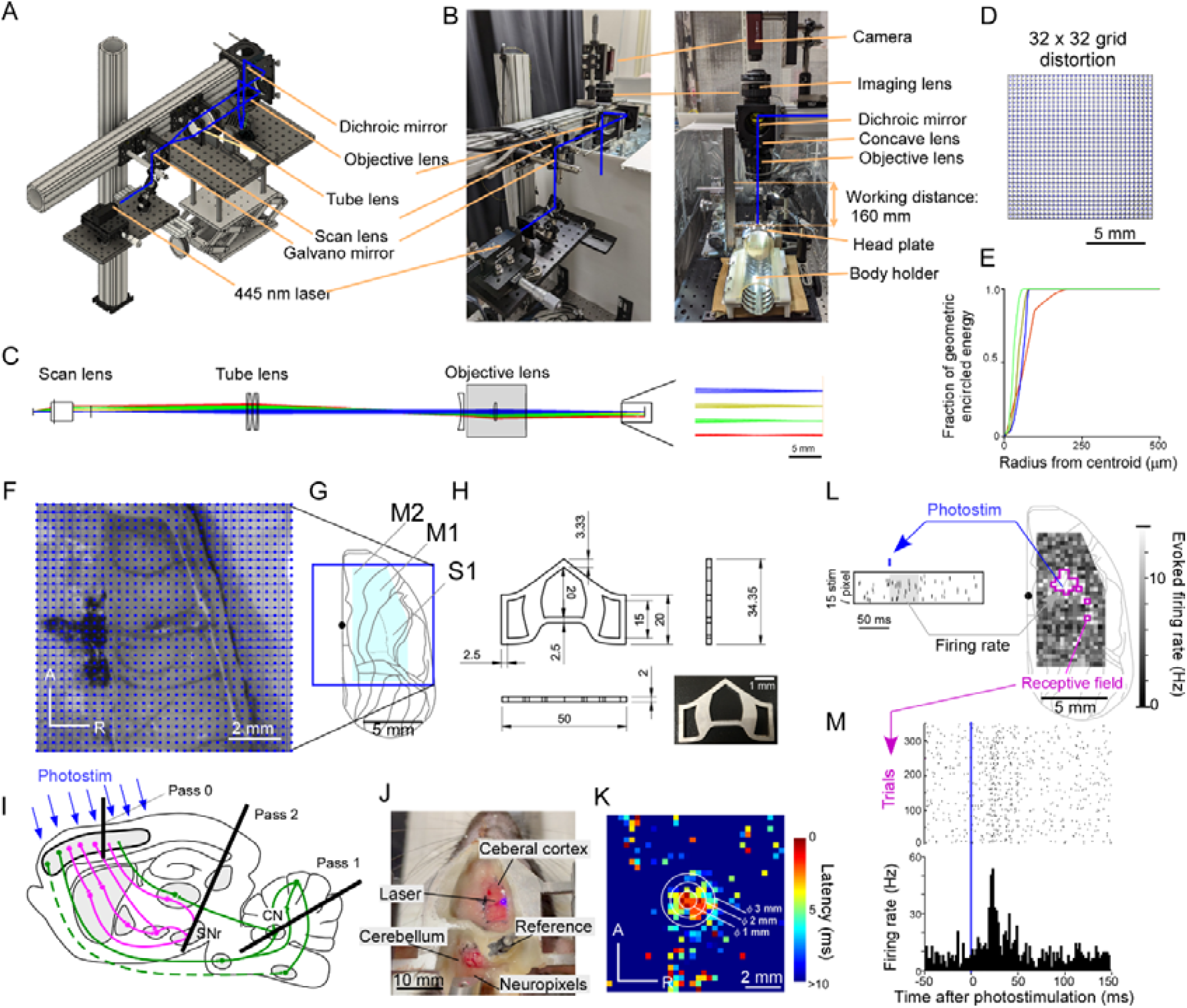
Optogenetic stimulation and electrophysiological recordings. **A**. 3D CAD data for the system. **B**. Photographs of the system. WD, working distance. **C**. Four laser beams correspond to scanning angles of 0, 4, 8, and 12 degrees, which focus at distances of 0, 2.4, 4.7, and 7.1 mm from the center, respectively. The objective lens (Nikon camera lens) was substituted by a lens with the same focal length. **D**. Grid distortion of 32 × 32 stimulation points (10 mm × 10 mm). **E**. Proportion of encircled energy as a function of the distance from the centroid. **F**. A photograph of thinned skull preparation. The blue dots correspond to the point of photostimulation. The blue vertical and horizontal lines indicate the intact skull bone. **G**. Dorsal view of brain atlas of right hemisphere. The blue square represents the 10 mm × 10 mm stimulation area. The black dot denotes the bregma. **H**. Headplate design. **I**. Approximate positions of recording pass 0 (cerebral cortex), pass 1 (for cerebellar nuclei (CN)), and recording pass 2 (for substantia nigra pars reticulata (SNr)). **J**. A photograph from the top during photostimulation of the cerebral cortex and recording at the cerebellum. **K**. Spike latency map of a single neuron in the cerebral cortex showing the resolution of photostimulation. **L** A representative responsive map for a single CN neuron. The purple polygons indicate the receptive field of the neuron. **M**. A raster plot (top) and PSTH (bottom) for trials in which photostimulation was performed on the receptive field in **L**.

### Histology

After all recording sessions, the rats were perfused transcardially with saline and subsequent 4% formaldehyde in 0.1 M phosphate buffer under deep anesthesia with urethane (3 g/kg, ip, Nacalai Tesque, Kyoto, Japan) The brains were removed and then post-fixed with the same fixative for at least 12 h at 4° Celsius. The brains were removed and post-fixed with the same fixative for >2 days. The brains were then embedded in 2 % agar in PBS and cut into sagittal sections with a thickness of 50 µm using a microslicer (VT1000S, Leica, Wetzlar, Germany). The serial sections were cover-slipped with mounting medium (DAPI Fluoromount-G, Southern Biotech, AL, USA). DiI, Venus, and DAPI (or NeuroTrace 435/455) were visualized through a fluorescence microscope (IX83 inverted microscope, Olympus, Tokyo, Japan).

### Tracer study

Adult Thy1.2-ChR2-Venus transgenic rats (n = 6, male: 2, female: 4, 180-400 g) were used in this experiment. The rats were anesthetized with 2.5% isoflurane and an air flow of 0.2 liter per minute. Injection of AAVretro-CAG-tdTomato (Tervo et al. 2016) (addgene, Plasmid #59462-AAVrg, 2.4×10^12^ GC/ml, 0.2 µL, 0.02 µL/min or 0.5 µL, 0.05 µL/min, 10 min) were performed (Hamilton, 10 µL Microliter Syringe Model #801 RN) at the VL (n=3, 1.7 mm left lateral to midline, 2.3 mm posterior to bregma, and 5.8-6.1mm deep from the brain surface), VM (n=2, 1.4 mm left lateral to midline, 2.4 mm posterior to bregma, and 6.9 mm deep from brain surface), or VA complex (n=1, 1.6 mm left lateral to midline, 1.7 mm posterior to bregma, and 6.0 mm deep from the brain surface) of thalamus by pressure through a glass micro pipette. Two weeks after AAV injection, the rats were anesthetized with urethane, and then perfused intracardially with saline and fixative solution containing 4 % paraformaldehyde. The Brain was quickly removed and fixed in 4% paraformaldehyde for 2 days at 4 °C then placed in a 30% sucrose in PBS solution for several days until they sank to the bottom of their container. The brain was then immediately immersed in isopentane over dry ice then stored at −80 °C until sectioned. Sagittal sections (20 μm or 50 μm) were obtained in a cryostat (NX50, Epredia) at −20*□* through the cerebellar nuclei, SNr and thalamus.

### Optogenetics

We simulated and assembled a scanning optical system with galvanometer mirrors (6210H, Cambridge Technology, USA) to systematically and rapidly photostimulate many areas of the cerebral cortex (**Fig. 1A-E**). Simulation of optical pathway was made with OpticStudio (version16.5, Zemax, USA) to illuminate an area of 10 mm × 10 mm, more than 150 mm away from the objective lens. This allowed us to put the electrode at the space between objective lens and the brain. After validation in the simulation environment, we built up scanning light pathway with SM2-based cage system (Thorlabs, USA) and the multifunction I/O device, USB-6259 (National Instruments, USA). Scan lens (EFL = 39 mm, LSM03-VIS, Thorlabs) and tube lens (two achromatic doublets (f = 400 mm, ACT508-400-A, Thorlabs) placed in a Plössl design) made an infinity-corrected laser light beam into the pupil of objective lens. We put a plano-concave lens (f = – 150 mm, LC1611-A, Thorlabs) just above the camera lens (Nikon, f=105 mm) to extend the working distance. We kept the diameter of the laser at pupil small enough (<0.1 mm) for achieving a large PSF extension in the Z-axis at each stimulus point. This minimized the light density shift due to the curvature of the rat cerebrum and at the same time reduce the light spreading conically inside the brain, thus creating a columnar illumination. For functional mapping of cerebral cortex, we stimulated 32 × 32 grid (323 μm spacing, total 1,024 points) with 115 ms interval pseudo-randomly for 15 cycles (Hira et al. 2009; Hira, Ohkubo, Tanaka, et al. 2013; Hira et al. 2015; Hira, Ohkubo, Ozawa, et al. 2013). The stimulation point after the thin-skull point was always at bone, so that the stimulation interval of cerebral cortex was more than 230 ms. For functional mapping of cerebellum, we stimulated 32 × 32 grid (129 μm spacing, total 1,024 points) with 230 ms interval pseudo-randomly for 15 cycles. Recording during the stimulation at the bone was monitored as an internal control to distinguish any contamination, including the neuronal activity responses to retinal light. We confirmed that there was no contamination of visually evoked activity in the responsive neurons that we analyzed. Each stimulation was 5 ms duration with 445 nm TTL-controllable diode laser (JUNO, Kyocera SOC, Japan). Laser power was set in each experiment sufficiently small not to induce movement when primary motor cortex was stimulated (0.7 mW–3.9 mW for stimulation of cerebrum, 15–21 mW for stimulation of cerebellum, at the center of field-of-view). Multi-unit activity (MUA) maps for varying laser power (**Fig. 2-1A-C)** indicated that the laser power we used was near the threshold at which evoked activity can be visualized. We also confirmed that the latency of the response and the shape of the map did not change significantly when we employed laser power that was strong enough to produce movement output. Even at the edge of the field of view (7.1 mm from the center), the laser power was >85.7% of the center. All system was controlled with custom-made software written with LabVIEW (National instruments, USA).

### Electrophysiology

We performed electrophysiological recordings following the methodology described in our previous report (Sugino et al. 2024). Right before the recording, the Neuropixels probes (Jun et al. 2017) (IMEC, Belgium) were manually coated with lipophilic dyes, DiI (DiIC18(3), PromoKine, Heidelberg, Germany). The probes were fixed to a holder and handled using a micromanipulator (SM-15M, Narishige, Japan). Reference and ground electrodes were soldered, and attached to a custom-made part with four independent silver wires located under the skull onto the cerebellum. The probe was lowered in the coronal plane at 5–10 degrees from vertical at a speed of 10 μm/s until 8 mm below pia for recordings of SNr, thalamus, striatum and cerebral cortex, and at 30–63 degrees from vertical at a speed of 10 μm/s until 8 mm below the dura for recordings of cerebellum. They were left to settle for more than 30 min to reduce subsequent drift due to post-insertion brain relaxation. Recording was controlled with Open Ephys software (version 0.5, https://github.com/open-ephys/plugin-GUI/releases) (Siegle et al. 2017). Spiking activity and local field potential (LFP) were obtained at 30,000 samples/s and 2,500 sample/s, respectively after 500 Hz high-pass filtering and band-pass filtering with 0.5 – 1kHz range, respectively. We recorded the timing and location of photostimulation at the same time with USB-6001 (National instruments, USA) to align timing of each photostimulation with the neural signals. When the Neuropixel probe was placed to penetrate the cerebellar nuclei, a “bird sign” appeared on the LFP correlation structure (**Fig. 2A**). Histological observation revealed that the presence of this bird mark corresponded to the CN. The wing of the bird would reflect the fact that similar LFP signal appeared from white matter around the CN so that the correlation between the white matters is high, while the CN has specific signal, reducing the correlation between CN and white matter. We finally determined location of each channel at the level of the subdivision within CN or SNr by histological observations.

**Figure 2.**
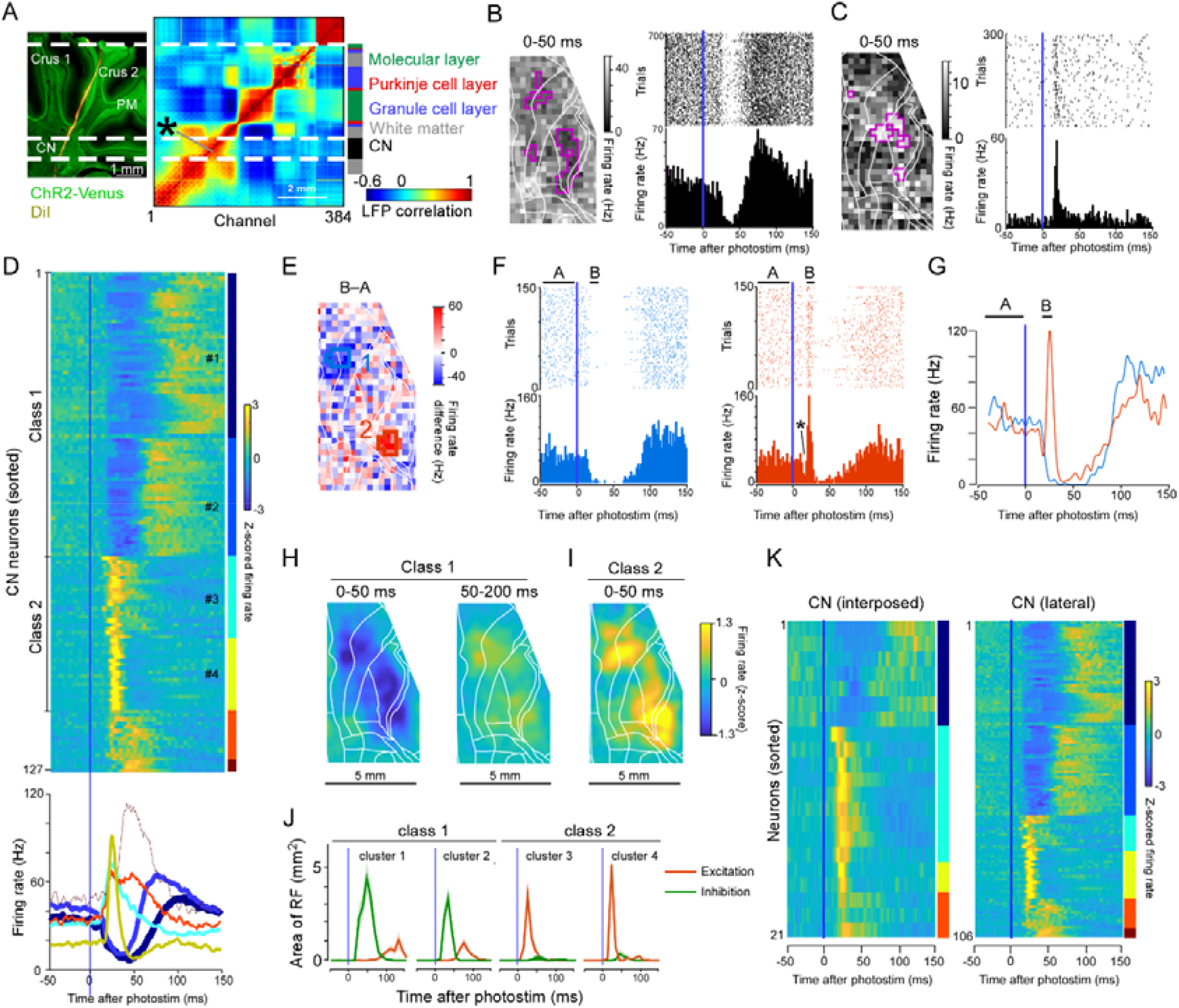
Two major classes of response patterns observed in cerebellar nuclei (CN) neurons. **A**. A sagittal section of cerebellum in a ChR2-Venus transgenic rat (left) and local field potential (LFP) correlation structure of the corresponding animal recorded with Neuropixels (right). DiI fluorescence shows electrode location. Dotted lines indicating boundaries of LFP correlation structure correspond to histologically confirmed cerebellar layer structure and cerebellar nuclei. **B**,**C**. Response maps of typical class 1 (**B**) and 2 (**C**) responses in CN (left). Magenta frames indicate responsive pixels. Raster plots and PSTHs for the responsive pixels (right). **D**. Z-scored PSTH of all CN neurons clustered by K-means clustering (top) and mean PSTH of each cluster (bottom). Colors correspond to cluster numbers. The thickness indicates the numbers of neurons in the cluster. **E**. A map of the difference in firing rate at two time bins: ™50–0 ms (A) and 20–30 ms (B) after stimulation of a CN neuron. **F**. Raster plots and PSTHs for the blue and red polygons in **E**. * indicates a transient inhibitory response before the fast excitatory response. **G**. An overlay of the PSTHs in **F. H**. Average input maps of class 1 neurons at 0–50 ms (left) and 50–200 ms (right) in CN. **I**. An average input map of a class 2 neuron in CN. **J**. Time course of excitatory and inhibitory responsive areas in clusters #1–4. K. The same analysis as **D**, but separated according to recording location. **Fig. 2-1** is available for more basic analysis.

**Figure 2-1.**
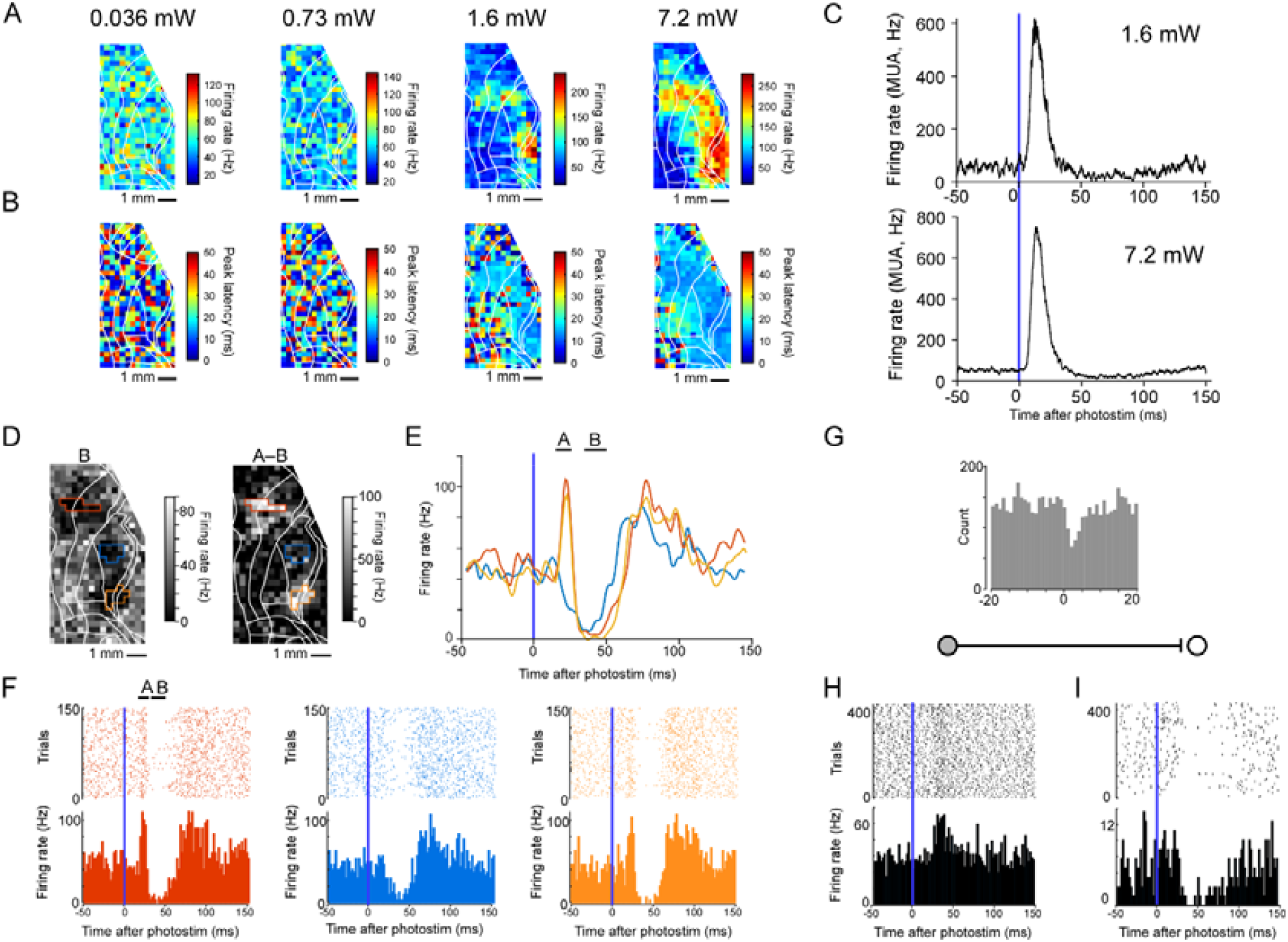
Cerebellar nuclei (CN) neuron responses. **A**. Laser power dependence of response maps of multi-unit activity in the cerebellar nucleus. **B**. Laser power dependence of response latencies in the same recordings as in panel **A. C**. PSTH of the same multi-unit activity as in A during stimulation at 1.6 mW (top) and 7.2 mW (bottom). **D**. Maps of firing rates 30–50 ms (B) after stimulation (left) and the difference in firing rates (right) at two-time bins: 20–30 ms (A) and 30–50 ms (B). **E**. Raster plots and PSTHs for the stimulus points enclosed by the red, blue, and orange polygons in **D. F**. Overlaid PSTHs. **G**. Cross-correlogram of simultaneous recordings of a PC (SS) and a neuron in CN. **H**. Rater plot and PSTH of the presynaptic PC (SS) in **G. I**. Rater plot and PSTH of the postsynaptic CN neuron in **G**. PSTHs in **H** and **I** were generated in trials with photostimulation of the receptive field of the PC for comparison.

We recorded neurons from lateral and interposed nucleus of CN. These regions are known to preferentially interact with the motor cortex (Harvey, Porter, and Rawson 1979; Mushiake and Strick 1993; Ohmae, Kunimatsu, and Tanaka 2017; Thach, Goodkin, and Keating 1992). We recorded neurons evenly from the SNr. We focused on the ML axis for the analysis because striato-SNr projection across AP axis seems to be similar (Lee, Wang, and Sabatini 2020), and both received the dorsomedial or dorsolateral striatal input where motor cortex sends the projections.

### Analysis of electrophysiological data

All analyses were performed with MATLAB (R2020a, Mathworks) or Python (version 3.7). Spike sorting was performed using KiloSort3 (https://github.com/MouseLand/Kilosort). Sorted clusters were manually curated with Phy template GUI (https://github.com/cortex-lab/phy). The spontaneous firing rate of each neuron was obtained by averaging the firing rates for 50 ms before photostimulation.

For the cerebellar recordings, we determined the cell types of the curated neurons in the following way. First, we identified each neuron’s region based on the LFP correlation structure described above. In the granule layer, we observed spiking activities with a NAW (negative after wave). Based on similar descriptions in rodents and monkeys (Beau et al. 2024; Muzzu et al. 2018; Prsa et al. 2009; Van Dijck et al. 2013; Van Kan, Gibson, and Houk 1993), we classified them as MFs. The complex spike of PCs was classified by a characteristic complex waveform and low firing rate (< 10 Hz) and the simple spike of PCs was identified according to recording location and high firing rate (> 20 Hz) (Sedaghat-Nejad et al. 2022; Sedaghat-Nejad et al. 2021; Cheron et al. 2009). We did not compare simple and complex spikes from the same neurons because we did not obtain sufficient numbers of PCs with both simple and complex spikes.

To identify “responsive neurons” to cerebral cortex stimulation and “receptive fields” in each responsive neuron, we first extracted the candidate points that satisfied the following inequalities:

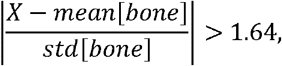

Where *X* is the mean firing rate of the neuron 0–50 ms after stimulation and [bone] indicates stimulation at the skull surface. Then, we excluded the candidate points surrounded with ≤ four candidate points among eight surrounding points. Finally, if any candidate point was surrounded by ≥ five candidate points among eight surrounding points, we defined the neuron and candidate points as responsive neuron and receptive fields, respectively. This criterion collects stimulated points that form a local cluster based on the fact that the distance between the points is smaller than the resolution of the stimulus. Among 10,000 randomly shuffled data, no data satisfied this criterion. We confirmed that it identified fewer neurons than those that were visually identifiable, which allowed us to limit our analysis to neurons that had responded unambiguously.

PSTHs were created by collecting the firing responses to the receptive fields from 50 ms before to 150 ms after photostimulation in each trial. The peak latency was obtained by finding a maximum or minimum value after convolving the PSTH with a Gaussian function with sigma of 3 ms. To examine onset latency, the value of convoluted PSTH at the time of stimulation and its maximum or minimum up to 50 ms after stimulation was standardized to 0 and 1, respectively. The time at which this value first exceeded 0.4 was determined to be the onset latency. To obtain the average responsive maps, we translated each map based on the position of bregma and angle of midline before averaging. K-means clustering was done by the MATLAB function “kmeans”. For each pair that was simultaneously recorded, a −50 ms to 50 ms cross-correlation range of spiking activity was calculated, and a significant interaction was defined when the peak was > 99.8% of the peak synchronization level calculated by shuffling spikes in the same window 1000 times. Among them, we visually inspected pair by pair to determine the putative monosynaptic interactions.

### Simulation of a spiking neural network

We created a spiking neural network with the Izhikevich model (Izhikevich 2003, 2007) in the Brian2 simulator in Python. The time step was set at 0.1 ms. The simulation time unit was ms. The Izhikevich model is described as follows:

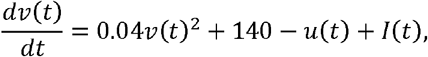

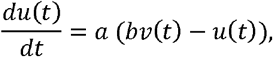

where *u* is potassium channel conductance, *u* is membrane potential, *a* is the leak current of the potassium channel, *b* is the voltage dependence of the potassium channel, *c* is post-depolarization potential, and *d* is the opening of the potassium channel after firing. When *v* exceeded 15 mV, the neuron was regarded to have an action potential, and *v* and *u* were reset to *c* and *u + d* respectively. We determined parameters *a,b,c*, and based on previous studies (Thibeault and Srinivasa 2013; Muddapu et al. 2019; Izhikevich 2007) (**Table 2**).

**Table 1.** A lens data list of the scanning optogenetics system. Shaded areas highlight substitutes of the Nikon camera lens with f = 105 mm.

| # |  | Type | Comment | Radius | Thickness | Material | Semi-Diameter | Mesh<br>Semi-Dia |
| --- | --- | --- | --- | --- | --- | --- | --- | --- |
| 0 | OBJECT | STANDARD |  | Infinity | Infinity |  | 0 | 0 |
| 1 | STANDARD | STANDARD |  | Infinity | 0 |  | 5.5 | 5.5 |
| 2 | STOP | STANDARD |  | Infinity | 0 |  | 0.5 | 0.5 |
| 3 |  | STANDARD | STANDARD | Infinity | 24.75 |  | 5.5 | 5.5 |
| 4 |  | STANDARD | STANDARD | Infinity | 4.25 |  | 10.761 | 10.761 |
| 5 |  | STANDARD | Mounting surface | Infinity | -4 |  | 17 | 17 |
| 6 | (aper) | BLACKBOX | LSM03-VIS.ZBB | Infinity | 29.935 |  | 12.7 | 12.7 |
| 7 |  | STANDARD | Last mesh<br>surface | Infinity | 25.15 |  | 8.621 | 8.621 |
| 8 |  | STANDARD |  | Infinity | 215 |  | 8.134 | 8.134 |
| 9 | (aper) | STANDARD |  | 760 | 2 | SF2 | 25.4 | 25.4 |
| 10 | (aper) | STANDARD |  | 186.8 | 5 | N-BK7 | 25.4 | 25.4 |
| 11 | (aper) | STANDARD | AC508-400-A | -219.8 | 0 |  | 25.4 | 25.4 |
| 12 | (aper) | STANDARD |  | Infinity | 2 |  | 15.7 | 15.7 |
| 13 | (aper) | STANDARD | AC508-400-A | 219.8 | 5 | N-BK7 | 25.4 | 25.4 |
| 14 | (aper) | STANDARD |  | -186.8 | 2 | SF2 | 25.4 | 25.4 |
| 15 | (aper) | STANDARD |  | -760 | 0 |  | 25.4 | 25.4 |
| 16 | (aper) | STANDARD |  | Infinity | 280 |  | 15.499 | 15.499 |
| 17 | (aper) | STANDARD | LC1611-A | -77.19 | 4 | N-BK7 | 25.4 | 25.4 |
| 18 | (aper) | STANDARD |  | Infinity | 0 |  | 25.4 | 25.4 |
| 19 | (aper) | STANDARD |  | Infinity | 41 |  | 11.193 | 11.193 |
| 20 | (aper) | STANDARD | KPX385 | 54.47 | 3.4 | BK7 | 12.5 | 12.5 |
| 21 | (aper) | STANDARD |  | Infinity | 41 |  | 12.5 | 12.5 |
| 22 | (aper) | STANDARD |  | Infinity | 0 |  | 13.982 | 13.982 |
| 23 | (aper) | STANDARD |  | Infinity | 163.483 |  | 25.4 | 25.4 |
| 24 | IMAGE | STANDARD |  | Infinity | 0 |  | 7.004 | 7.004 |

**Table 2.** The numbers of neurons, numbers of domains, spontaneous firing rate, and four parameters of the Izhikevich model. FSI: fast spiking interneuron, MSN: medium spiny neuron, GPe: external segment of global pallidus, STN: subthalamic nucleus, SNr: substantia nigra pars reticulata, GPi: internal segment of global pallidus, ST: stellate cell, BA: basket cell, GO: Golgi cell, GR: granule cell, PC: Purkinje cell, IO: inferior olive cell, MF: mossy fiber, DCN: deep cerebellar nucleus, VM: ventromedial nucleus, VL: ventrolateral nucleus. L5PN; layer 5 pyramidal neuron. The number of neurons in each type in basal ganglia, cerebellum, and PF were determined by the single-cell-resolution whole-brain atlas (Murakami et al. 2018).

| Region | Cell type | No. of neurons | No. of domains | Spont. firing rate (Hz) | a | b | c | d |
| --- | --- | --- | --- | --- | --- | --- | --- | --- |
| Basal ganglia | FSI | 3200 | 32 | 28 (Berke et al. 2004; Gage et al. 2010) | 0.1 | 0.2 | -65 | 2 |
| Basal ganglia | MSN | 32000 | 32 | 7 (Miller et al. 2008) | 0.02 | 0.2 | -65 | 8 |
| Basal ganglia | GPe | 18000 | 36 | 24 (Mallet et al. 2008) | 0.02 | 0.2 | -65 | 8 |
| Basal ganglia | STN | 3600 | 6 | 14 (Mallet et al. 2008; Yasoshima et al. 2005) | 0.02 | 0.2 | -65 | 8 |
| Basal ganglia | SNr / GPi | 12600 | 14 | 32 (Maurice et al. 2003) | 0.02 | 0.2 | -65 | 8 |
| Cerebellum | ST | 3510 | 1 | 20 (Jelitali et al. 2016) | 0.1 | 0.2 | -65 | 2 |
| Cerebellum | BA | 910 | 1 | 20 (Jelitali et al. 2016) | 0.1 | 0.2 | -65 | 2 |
| Cerebellum | GO | 650 | 1 | 18 (Muzzu et al. 2018) | 0.1 | 0.2 | -65 | 2 |
| Cerebellum | GR | 10000 | 1 | 5 (Jelitali et al. 2016) | 0.2 | 0.2 | -65 | 0.05 |
| Cerebellum | PC | 1040 | 1 | 68 (Muzzu et al. 2018; Jelitali et al. 2016) | 0.02 | 0.2 | -65 | 0.01 |
| Cerebellum | IO | 130 | 1 | 1 (Jelitali et al. 2016) | 0.1 | 0.2 | -65 | 2 |
| Cerebellum | MF | 1170 | 1 | 15 (Muzzu et al. 2018) | 0.2 | 0.2 | -65 | 0.05 |
| Cerebellum | DCN | 2150 | 2 | 50 (this study) | 0.02 | 0.25 | -65 | 0.05 |
| Thalamus | PF | 7200 | 6 | 8 (Urbain et al. 2015; Marlinski and Beloozerova 2014) | 0.02 | 0.25 | -65 | 0.05 |
| Thalamus | VM | 200 | 1 | 8 (Urbain et al. 2015; Marlinski and Beloozerova 2014) | 0.02 | 0.25 | -65 | 0.05 |
| Thalamus | VL | 200 | 1 | 8 (Urbain et al. 2015; Marlinski and Beloozerova 2014) | 0.02 | 0.25 | -65 | 0.05 |
| Cerebral cortex | L5PN | 200 | 1 | 12 (O'Connor et al. 2010) | 0.02 | 0.2 | -65 | 8 |

We set the network connectivity of basal ganglia in detail based on recent anatomical studies at the submodule level (Foster et al. 2021; Jeon et al. 2022; Hintiryan et al. 2016) (**Fig. 7-1**). Conversely, although recent studies have provided essential knowledge (Pisano et al. 2021; Kebschull et al. 2020; Wu et al. 2023), because of limited projectome data of the cerebro-cerebellar loop circuit at a subdivision level (for instance, at the level of the microzone (Sugihara et al. 2009)), we modeled only a part of the cerebellar network with known anatomical connections. The synaptic circuit between submodules was shown in **Fig. 7-1A**. The connection probability (*p*) was a value between 0.008 and 0.15 and was assumed to be random. This probability was determined from previous studies and the axon density and was determined to match the results of our electrophysiology experiments. The connection probability where there is no anatomical connection was zero. There were no recurrent connections between the same submodules except those explicitly shown in **Fig. 7-1A**. We set the conduction delay of each connection as shown in **Fig. 7-1**. The delay was the same across the same connection types, but we varied the delay in three parts of the cerebellar circuit, i.e. MF → Granule cell, PF → CN, and Granule cell → PC. The variations in the delay of the three connections were defined as follows.

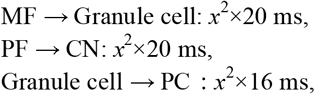

where *x* was a random value between 0 and 1 for each pair of neurons. Each synaptic input was delivered immediately after the delay time as a transient voltage step in the Izhikevich model. Conduction delay and synaptic weight parameters were determined in agreement with the results of our electrophysiological experiments based on previous studies. The simplified diagram combining all the simulated networks with connection probability, conduction delay, and synaptic weight is shown in **Fig. 7A**.

For simplicity, we did not consider the following aspects in our model: gap junction, specific waveform of complex spike, pose of simple spike after the complex spike, short-term synaptic plasticity, long-term synaptic plasticity of most neuron pairs, dendritic structure, dendritic spike, neuromodulator, glial cells, and vascular endothelium.

As an input from the cerebral cortex, we simulated a Poisson spike train as a spiking activity of association cortex (N = 600 neurons). The spontaneous firing rate was set to 10 Hz, and the firing rate after virtual optogenetic stimulation was set to 100 Hz (5 ms). Virtual optogenetic stimulation was administered 15 times during 3000 ms of stimulation at various intervals. By visualizing a PSTH of each subdivision or each region, we manually modified synaptic weight, connection probability, and conduction delay by referring to electrophysiological data obtained in this study. Note that although we searched a broad set of parameters, the overall connection diagram was not changed, indicating that the circuit is consistent with our anatomical knowledge. After determining the parameters, we ran the simulation for longer time periods (6000 ms including 20-30 stimulations), which were repeated 5 times.

### Reservoir based reinforcement learning model

The dynamics of the cerebral cortex were modeled as a reservoir circuit described as

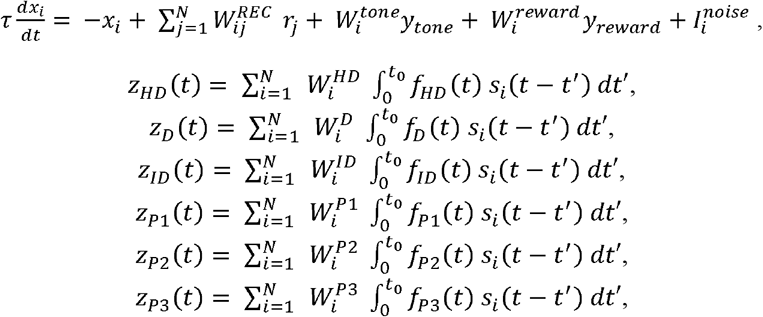

where *r*_*i*_ = *tanh(x*_*i*_) represents the firing rate of the *i-th* recurrent unit (*i* = 1,…,*N*), *S*_*i*_ = (*r*_*i*_+1)^2^, and the filter functions *f*_*HD*_, *f*_*D*_, *f*_*ID*_, *f*_*P*1_, *f*_*P*2_, *f*_*P*3_ were defined by the double-exponential function fitted to the experimental data (**Fig. 8B**). Because output neurons in the cortex are excitatory, *S*_*i*_ limits the output of the reservoir to positive values. It can also mimic the distribution of the firing activity of cortical neurons by converting it to highly skewed activity. The reservoir network was simulated at the time step of 1 ms. The variable *y*_tone_ and *y*_reward_ represents the input of tone and reward, respectively (1 or 0). *N*=800 is the number of units, and τ= 20 *ms* is the unit time constant. Since many cerebral cortices, including posterior parietal cortex, secondary motor cortex, and sensory cortex in rodents have neurons that respond to tone cue and reward signals (Hira et al. 2023; Barthas and Kwan 2017; Lacefield et al. 2019), both were added to the reservoir as inputs. The recurrent connectivity is represented by the *N × N* sparce and Kwan matrix 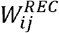 where the connection probability between units is *p* = 0.14, and the weight follows a Gaussian distribution with a mean of zero and a standard deviation of 0.189. The activity of the network was read out by *z*_*HD*_, *z*_*D*_, *z*_*ID*_, *z*_*Pl*_, *z*_*P2*_, and *z*_*P3*_ through the output connectivity matrix 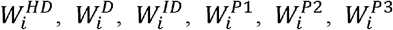, with initial values determined by a Gaussian distribution with a mean of zero and a standard deviation of. Here, because we regarded phase 3 as a rebound of phase 2, 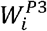 was the same as 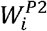. The input weight vectors 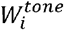 and 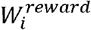 followed a Gaussian distribution with a mean of zero and a standard deviation of The values of *y*_*tone*_ and *y*_*reward*_ were 0, except during a tone or reward, comprising a step with a duration of 50 ms and amplitude of 1. A noise current was included as a *N × 1* random vector, 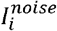 defined with a Gaussian distribution with a mean of zero and a standard deviation of 0.001. Reservoir circuits were designed to mimic the complex dynamics of the cortex, not to optimally configure the state for reinforcement learning.

Our spiking neural network is at semi-realistic scale and has a relatively large number of neurons (2.4×10^5 total neurons). It took about 20 min, or about 400 times longer to run the 3 sec simulation (Mother board: Z690/D4, CPU Intel Core i9-12900K 16 cores 3.2GHz, RAM 128GB). If this were used directly as a reinforcement learning model, it would take 13,300 hours to simulate 100 iterations in 20 minutes, which is practically impossible. Therefore, the cortex was modeled as a reservoir network, and the outputs of the basal ganglia and cerebellum were modeled as linear filters based on experimental data. Note that it took about 98 minutes for 100 iterations of the 20-min reinforcement learning model in our environment. Under these conditions, 123 hours were needed to obtain the data in Fig. 8J. Maass and colleagues (Chen, Scherr, and Maass 2022) have proposed a fast method for including plasticity at a scale comparable to that of our semi-realistic spiking neural network. They have successfully imposed learning equivalent to backprop on a large spiking neural network. We believe that semi-realistic spiking neural networks could be used to elucidate the learning mechanism of our model in the future by using an appropriate learning rule.

### Reinforcement learning task

The agent received the tone cue and reward as inputs and determined whether to hold or pull the lever at each time step of 100 ms. When the agent held the lever for 1 s, the tone cue was presented. If the agent pulled the lever before holding the lever for 1s, the tone cue was not presented, and the agent needed to hold the lever for 1s for the next tone cue. The agent needed to pull the lever within

0.3 s after tone cue presentation to get the reward. If the agent did not pull the lever within 0.3 s, the agent did not get the reward and needed to hold the lever for 1 s to receive the next tone cue. We have shown that rats can learn this task quickly (within a day of training) (Kimura et al. 2012; Kimura et al. 2017; Kawabata et al. 2020; Yoshizawa et al. 2023; Saiki et al. 2018).

The agent determined whether to pull or hold the lever every 100 ms, by comparing the pull value (*q*_*pull*_) and hold value (*q*_*hold*_). We defined these values as follows:

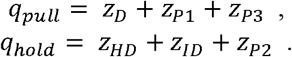

Here, the excitatory and inhibitory effect on the thalamus was regarded as the pull value and hold value, respectively. If the sum of these activities integrated in M1 from the thalamus or in the brainstem circuits exceeded the motor threshold, the lever was pulled. Thus, with *q*_*pull*_ and *q*_*hold*_, we defined the probability of pulling, *q*_*pull*_, as follows:

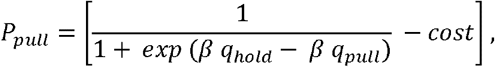

where [*x*] returns *x* if *x* is positive and 0 if is zero or negative, *β* = 1.2 is inverse temperature, and cost = 0.45 is the bias parameter. Every 100 ms, the maximum value of this probability was calculated, and this value determined pull or hold. A pull at the appropriate time was rewarded; however, there was a time lag of 0.2 s between the decision to output the action and the reward. This time lag was broadly consistent with the experimental data.

### Learning

This simple model assumed that synaptic plasticity does not occur within the reservoir but only affects the direct pathways and PFs. All the synaptic plasticity was based on RPE, *δ*_*t*_, defined as follows:

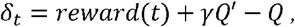

where the discount rate *γ* was set to a half-life of 3 s. *Q* and *Q’* are values of consecutive time steps (Note that the time step of learning is 100 ms) as follows:

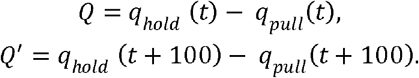

Synaptic weight was renewed every 1 s (10 behavioral steps). Under the default condition, synaptic changes were made to the direct pathway and phase 2 as follows:

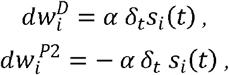

where the learning rate was *α* = 0.0025. This difference in timescale was employed because the timescale in which LTP actually occurs is slower than the timescale for coincidence detection at the synapse. Additionally, random synaptic fluctuations with a standard deviation of 0.001 were added, and each synaptic weight was divided by the sum of synaptic weights in the corresponding pathways according to homeostatic plasticity. As phase 3 activity in the cerebellar nuclei was likely to be a rebound of phase 2 activity, 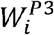 was changed in conjunction with 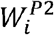.

Synaptic changes in the direct pathway and PF-PC pathway in the cerebellum correspond to the dopamine-dependent LTP in the corticostriatal synapse (Yagishita et al. 2014) and CF-dependent LTD in the PF-PC synapse, respectively, when RPE is positive. Negative RPE increases local ACh concentration (Chantranupong et al. 2023), and cholinergic input induces LTD in the D1-positive medium spiny neuron (Shen et al. 2015). PF-PC synapses undergo LTP when PCs only exhibit simple spikes without complex spikes (Hirano 1990). Accordingly, we allow for bidirectional plasticity in both direct and PF-PC pathways.

To investigate how this learning depends on the properties of this model, we ran the same simulations with different parameters as follows. For each of the six unit time constants of 5, 10, 20, 30, 40, and 50 ms, we ran with delays of 0, 10, 20, 30, 40, 50, 60, and 70 ms on the basal ganglia or cerebellar signals. For noise dependence, we ran it by adding Gaussian noise with standard deviations of 0, 1, 2, 5, 10, and 20 times the standard deviation of the reservoir’s output signal. When examining oscillation dependence, the Unit time constant was fixed at 50 ms. For the oscillation frequency dependence, simulations were performed by adding sine waves of 10, 15, 20, 25, 30, 35, 40, 60, and 100 Hz (the magnitude of the oscillation was 1.25 times the standard deviation of the input current) to the input current to the Unit. For the magnitude of oscillations, simulations were performed with 20 Hz sine waves of magnitudes 0.1, 0.2, 0.5, 0.75, 1, 1.25, 1.5, 1.75, 2.5, and 4.2 times the standard deviation of the input current. For the delay time, 20 Hz sine waves (1.25 times the standard deviation of the input current for the magnitude of the oscillation) were run with delays of 0, 10, 20, 30, 40, 50, 60, 70, 80, and 90 ms for the signal in thein the basal ganglia or cerebellum.

The same simulation was repeated with the time step for action and value updates changed from 100 ms to 50 ms (**Fig. 8-1L-N**). The probability of pulling the lever was set to 1/2, as 50 ms doubles the number of times per a decision has to be made. The discount rate was set to the square root of the original value due to the change in bin width. The learning rate, amplitude of reward, and inverse temperature were set to the same value because the amount of reward per unit time did not change.

### Statistics

The Wilcoxon’s rank-sum test and *χ*^2^-test were used for pairwise comparisons. Data were expressed as mean□ ± □ S.E.M in the experimental data, ± □ S.E.D. in the simulation data unless noted otherwise.

## Results

### Transcranial optogenetic stimulation of cerebral cortex and neuronal recordings in CN and SNr

To stimulate large cortical areas in a point-by-point manner and to make comparisons between stimulated areas, we used transgenic rats (Mitani et al. 2022; Saiki et al. 2018) densely expressing ChR2 under the Thy1.2 promoter in all layers of the cerebral cortex under head-fixed conditions in the awake state. We previously reported a method enabling photostimulation of ChR2-expressing neurons in mice with retained skulls (TOS, transcranial optogenetic stimulation) (Hira et al. 2009; Hira, Ohkubo, Ozawa, et al. 2013; Hira et al. 2015; Hira, Ohkubo, Tanaka, et al. 2013). For optical stimulation of the rat cerebral cortex, the area that can be stimulated must be expanded. Therefore, we simulated the optical pathway and constructed a system with a photostimulation area of 10 mm × 10 mm based on the simulation results (**Fig. 1A–E**). We confirmed that photostimulation with our new system was possible in the transgenic rats by maintaining the transparency of a thinned skull in one hemisphere of the dorsal cerebral cortex (**Fig. 1F–J**). To determine the resolution of photostimulation, we stimulated neurons in the cerebral cortex with a 445 nm laser and recorded the surrounding spiking activity. Spiking responses of < 2 ms latency were observed in neurons < 0.5 mm from the center of optical stimulation in the cerebral cortex (**Fig. 1K**), indicating that direct photostimulation occurred within 0.5 mm from the center of the photostimulation site.

We divided a 10 mm × 10 mm area including a thinned-skull portion into 32 × 32 points and illuminated each point with a focused blue laser for 5 ms (**Fig. 1F**). We recorded CN and SNr neural activity using Neuropixels probes (Jun et al. 2017). In total, 544 and 282 neurons were isolated in CN (9 rats, 23 sessions) and SNr (8 rats, 19 sessions), respectively. We examined the exact recording location of each channel with histological data and local field potential (LFP) correlation structures. We identified the location of all recorded neurons at the level of subdivision of each region or layer (interposed or lateral nuclei for CN, molecular layer, granule cell layer, PC layer, white matter for neurons in the cerebellar cortex, medial (< 2mm from midline) or lateral (≥2 mm from midline) portion for SNr,).

Responses of a representative neuron recorded at lateral CN are shown in **Fig. 1L,M**. We displayed the average firing rate between 0–50 ms from the stimulus at each point on a color scale (hereafter referred to as the response map, **Fig. 1L**). The response map indicated that the neuron received the input from the primary sensory cortex (S1), primary motor cortex (M1), and secondary motor cortex (M2). We called the stimulation points that significantly affected the firing rates of recorded neurons the “receptive field” of the neuron. A raster plot and peri-stimulus time histogram (PSTH) for the receptive field are shown in **Fig. 1M**. We performed this analysis on all neurons and identified responsive neurons that had receptive fields.

### Two types of response classes in CN neurons

To identify CN neurons, DiI fluorescence dye was applied to the Neuropixels probes, and the probe trajectories were identified histologically (**Fig. 2A**). We used 384 electrodes every 20 mm to skip one recording site, so that the recordings were linear over 7.5 mm. LFP correlation analysis revealed multiple clusters, which corresponded well with the histological layer structure and nuclear boundaries. In particular, we found the bird sign of the LFP (**\*** in **Fig. 2A**) enabling us to separate CN from their surroundings (see Methods). This corresponds to the fact that CN are located in white matter and are highly correlated with signals from white matter.

We identified two representative CN response classes. A typical class 1 response was a strong inhibitory response followed by an excitatory response (**Fig. 2B**), whereas a typical class 2 response was transient excitation (**Fig. 2C**). K-means clustering (K = 6) of z-scored temporal response patterns of all responsive CN captured these two clusters: clusters #1 and #2 for class 1, and clusters #3 and #4 for class 2 (**Fig. 2D**). In clusters #1 and #2 (class 1), the onset and peak latency of inhibitory responses were 13.1 ± 0.6 ms and 41.2 ± 3.3 ms, respectively, and the onset and peak latency of excitatory responses were 58.9 ± 3.2 ms and 94.6 ± 2.9 ms, respectively. Neurons with class 1 response accounted for 56.7% (72/127) of all neurons. In clusters #3 and #4 (class 2), the onset and peak latency of excitatory responses were 14.0 ± 0.7 ms and 23.9 ± 0.4 ms, respectively. The excitatory peak of class 2 responses was significantly faster than that of class 1 inhibitory responses (p = 2.9 ×10^−12^, Wilcoxon’s rank-sum test). Neuron with class 2 response accounted for 30.7% (39/127) of all neurons. Thus, neurons with class 1 and class 2 responses cover the majority (87.4%, 111/127) of CN responsive neurons. Neuron with class 1 response had a higher spontaneous firing rate than the neuron with of class 2 response (37.9 ± 2.0 Hz vs. 27.6 ± 4.0 Hz, p = 0.0053, Wilcoxon’s rank-sum test), suggesting that neurons with class 1 and 2 responses have distinct biophysical properties.

It is unclear whether neurons with class 1 and class 2 responses are two completely different groups of neurons or whether a single neuron has both properties. Notably, some neurons exhibited mixed responses of class 1 and class 2. A neuron that was classified as class 1 had a transient fast excitatory response resembling class 2 only when S1 was stimulated (**Fig. 2E–G**). In other words, the class 2 response pattern was added only in S1, on top of the inhibitory to excitatory response pattern typically observed in class 1. Another neuron that was classified as class 1 had a class 2-like fast transient excitatory response when M2 or S1 was stimulated (**Fig. 2-1D–F**). Thus, class 1 and 2 responses can merge within a single neuron. Distinct response patterns depending on the stimulation areas indicate that CN can integrate the activity of multiple cortical areas in a manner that reflects area-specific time series information.

Furthermore, it is unclear whether the different response patterns of class 1 and class 2 depended on the location of stimulation or recording. The class 1 excitatory and inhibitory responses and the class 2 excitatory responses were evoked by a wide area of M2, M1, and S1, with no clear differences (**Fig. 2H**). The receptive fields of class 1 inhibitory and class 2 excitatory responses had receptive fields of up to 4–5 mm^2^, with no clear differences (**Fig. 2I**). By contrast, more class 1 responses were observed in the lateral nucleus than in the interposed nucleus (p = 3.8 × 10^−3^, χ^2^-test), whereas more class 2 responses were observed in the interposed nucleus than in the lateral nucleus (p = 0.040, χ^2^-test; **Fig. 2K**). Thus, CN response patterns depend more on the location of the recording than that of the stimulus.

### Recordings of PCs and MFs explain responses of CN neurons

There are multiple possible pathways from the cerebral cortex to the CN, and the pathways through which neurons with class 1 and 2 responses are generated remain unclear. We measured the activity of two structures that enter CN: MFs and PCs. MFs exhibit characteristic short waveforms of axonal activity and negative after waves (NAW) (Beau et al. 2024; Muzzu et al. 2018; Prsa et al. 2009; Van Dijck et al. 2013; Van Kan, Gibson, and Houk 1993). We determined MFs by waveform features and the observed layer. Most MFs showed very fast transient excitatory responses to cortical stimulation (**Fig. 3A,B**). Inhibitory responses were observed in five MFs, whereas 87 MF excitatory responses had an onset and peak latency of 11.0 ± 0.5 ms and 21.4 ± 0.9 ms, respectively, which were significantly faster (by 3 ms and 2.5 ms) than those of class 2 CN (onset: p = 7.0 × 10^−6^; peak: p = 3.0 × 10^−5^, Wilcoxon’s rank-sum test; **Fig. 3G**). These results are consistent with the idea that fast excitatory responses of class 2 CN were via MF input, which is also anatomically plausible.

**Figure 3.**
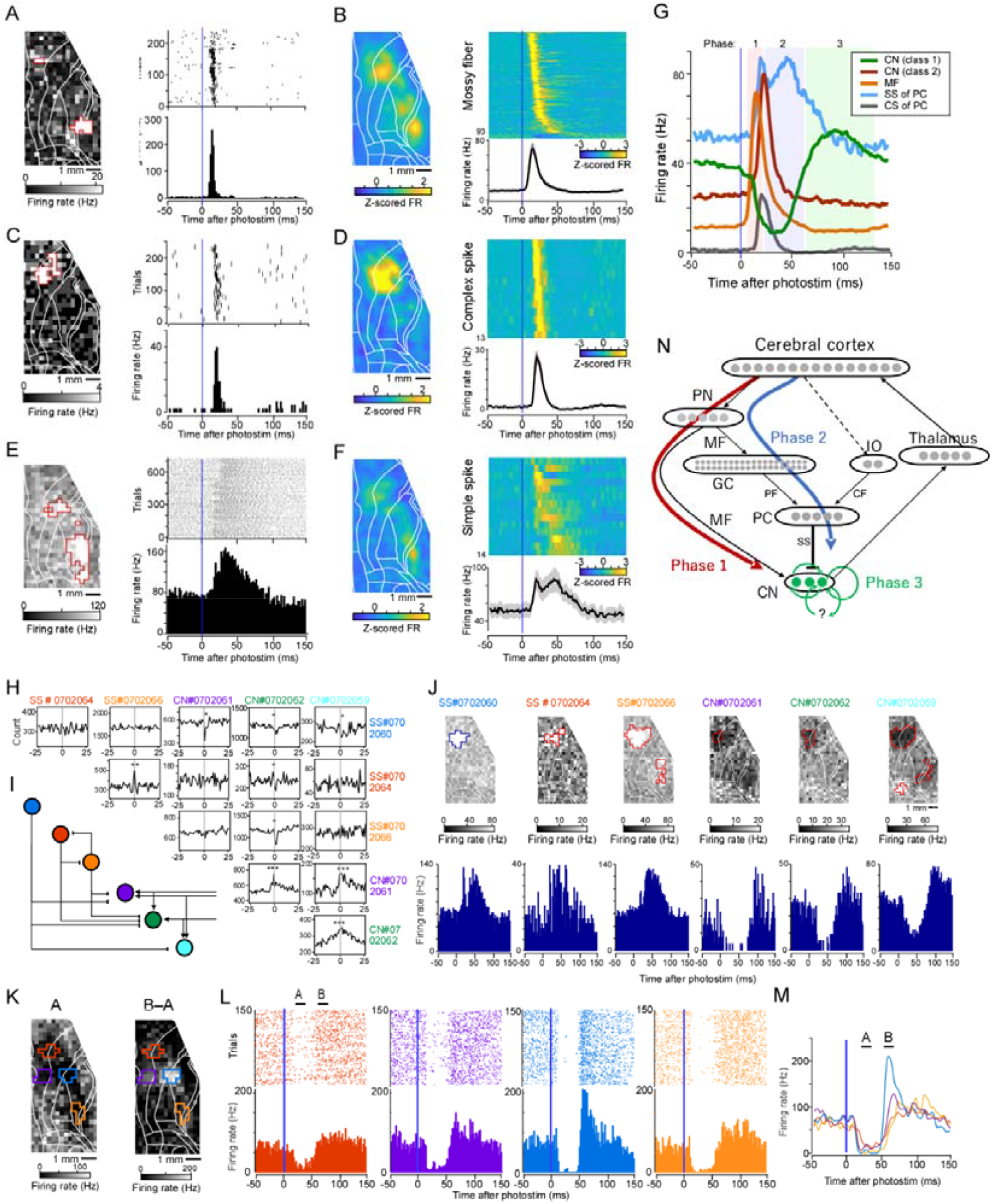
Activity propagation from the cerebral cortex to cerebellar nuclei (CN) neurons. **A**,**C**,**E**. A response map, raster plot, and PSTH of a mossy fiber (MF) neuron (**A**) and complex spike (**C**) and simple spike (**E**) of a PC. **B**,**D**,**F**. Normalized average responses (left) of MF neurons (**B**) and complex spikes (**D**) and simple spikes (**F**) of PCs. The order was sorted by the peak time. Average maps of all activities (right). **G**. Mean firing rates of CN neurons with class 1 responses, CN neurons with class 2 responses, MFs, SS (simple spike), and CS (complex spike). Shaded areas indicate phases 1–3. **H**. Cross-correlogram of simultaneous recordings of three PCs (SS) and three neurons in CN. **I**. A predicted synaptic circuit among neurons in **H**. J. Response maps and PSTHs of six neurons shown in **H**. All PSTHs were generated in trials with photostimulation of the receptive field of the blue neuron (SS#0702060) for comparison. **K**. Firing rates 20–40 ms after photostimulation (A in **L**) of a CN neuron (left) are shown as a map. Firing rates 50–70 ms after photostimulation (B in **L**) minus that of A of the same neuron are shown as a map (right). **L**. Raster plots and PSTHs of CN neurons from the color-corresponding areas specified in **K. M**. Mean firing rates of CN neurons from the color-corresponding areas in **K** when photostimulated. **N**. Possible pathways from the cerebral cortex to CN. Phases 1, 2, and 3 correspond to the time for excitatory input in class 1 response, inhibitory input in class 2 response, and excitatory input in class 2 response.

PCs are characterized by simple spike (SS) and complex spike (CS). We analyzed these two types of spiking activity separately. All complex spikes showed transient activity with jitter in a very narrow range (**Fig. 3C,D**; onset latency, 15.0 ± 0.6 ms, range, 12–19 ms; peak latency, 23.4 ± 0.7 ms, range, 20–28 ms). Notably, the cortical areas that elicited complex activity were confined to a portion of frontal cortex (**Fig. 3D**, left), and S1 rarely evoked complex spikes. SSs showed broad responses compared with complex spikes (**Fig. 3E,F**). The onset and peak latency were 15.4 ± 1.7 ms and 35.3 ± 3.1 ms, respectively. These values are comparable to those of class 1 inhibitory responses of CN. Additionally, the duration of the excitatory responses was roughly similar to that of CN class 1 inhibitory responses. Interestingly, the mean SS response had a transient decrease of firing activity 20–46 ms after photostimulation. This is compatible if the CS, which has a peak latency of 23.4 ms, pauses the SS for approximately 20 ms. We also examined the cross-correlograms of three putative SSs and three CN neurons in simultaneous recordings and found monosynaptic inhibition from SSs of PCs to CN neurons with class 1 response (**Fig. 3H,I**). Furthermore, the maps of inhibitory responses of CN and the maps of SS excitation overlapped (**Fig. 3J**). Similar result was observed in the other rat (**Fig. 2-1G-I**). Thus, monosynaptic inhibition from SS to CN neuron with class 1 response may directly contribute to the strong suppression of CN neurons with class 1 response after photostimulation. We examined all responsive MF-CN pairs closely and found no significant MF-CF pairs in cross-correlograms. We also closely investigated activity propagation between complex spikes and CN pairs, the equivalent of nuclear-olivary connections, and found no such pairs. To investigate the propagation of information between these pairs, it will be necessary to use multi-shank electrodes with higher electrode density in the future.

As the broad excitation after inhibition in class 1 response was not caused by MFs or PCs, it was most likely due to rebound excitation (Hoebeek et al. 2010; Aizenman and Linden 1999). By contrast, we found a CN neuron whose excitation after inhibitory activity differed across cortical areas (**Fig. 3K–M**). This neuron exhibited an increased excitatory response after photostimulation of M1 (blue, purple) but not M2 (red) or S1 (orange). Thus, the responses can switch in a region-specific manner. In summary, class 1 excitatory responses appear to be a rebound; however, the underlying process is complex, and the cerebral cortex may use this excitation to transmit region-specific information. Taken together, we illustrated the circuitry for phase 1 (excitation of class 2 responses), phase 2 (inhibition of class 1 responses), and phase 3 (excitation of class 1 responses) (**Fig. 3N**).

### Two types of response classes in SNr neurons

Next, we examined the pattern of transmission from the cerebral cortex to the SNr, an output nucleus of the basal ganglia. We histologically confirmed the recording location (**Fig. 4A**). We identified two types of response classes in SNr neurons. Typical neurons with class 1 responses showed a clear triphasic pattern similar to that of previous studies (**Fig. 4B**) (Fujimoto and Kita 1992; Nambu et al. 2000; Nambu, Tokuno, and Takada 2002; Sano et al. 2013; Kitano, Tanibuchi, and Jinnai 1998). The triphasic response consisted of an excitatory-inhibitory-excitatory activity sequence in which the third excitatory response was always larger than the first one, and the firing activity was often completely suppressed during the second inhibitory phase. We also found a monophasic excitatory response (class 2) (**Fig. 4C**). K-means clustering (K = 6) of z-scored temporal response patterns of all responsive SNr neurons roughly captured these two classes: clusters #1 and #2 for class 1, and clusters #3 and #4 for class 2 (**Fig. 4D**). Neurons with class 1 (24/56) and class 2 (25/56) responses cover the majority (87.5%, 49/56) of SNr neurons.

**Figure 4.**
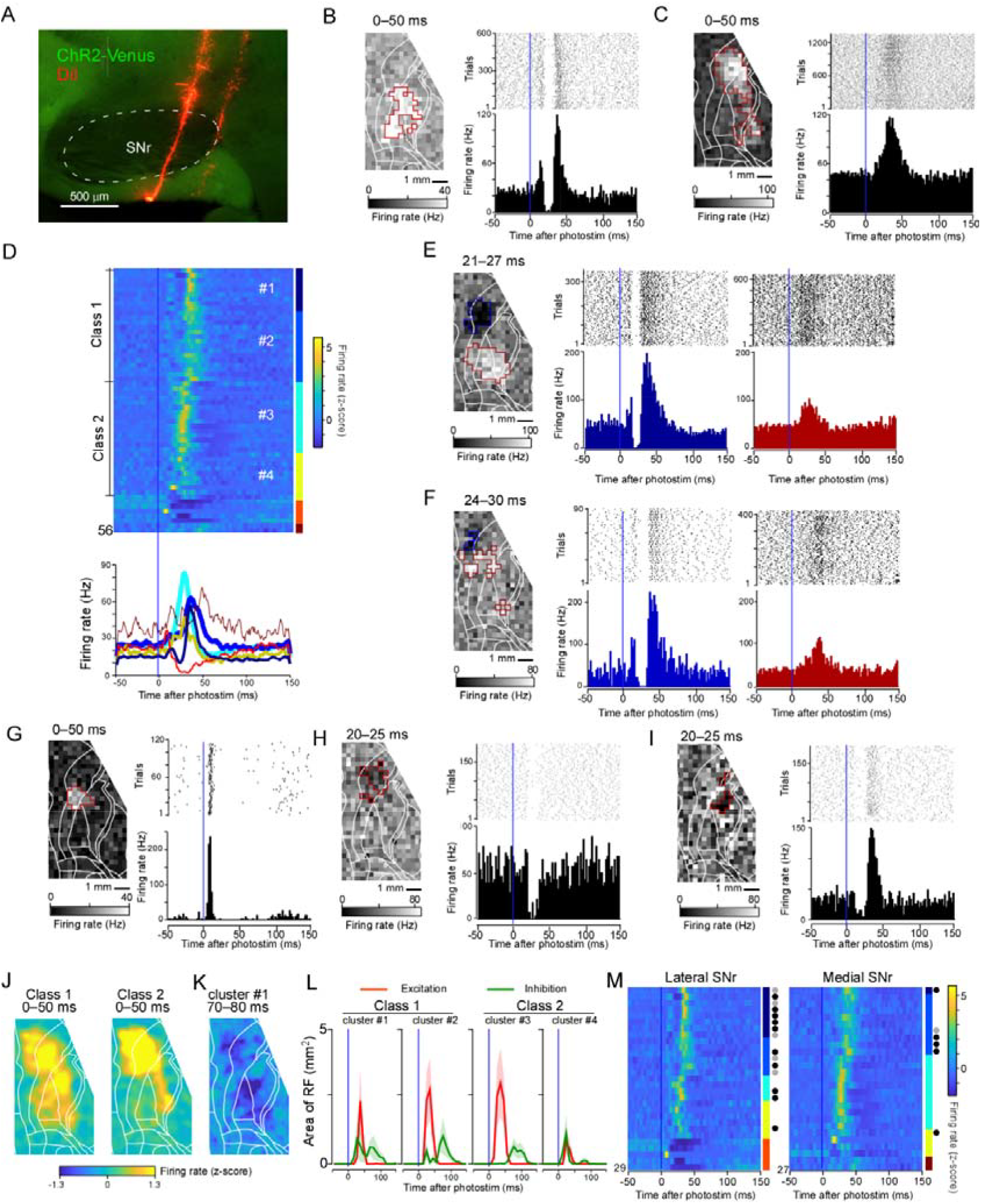
Analysis of single neuron response patterns of substantia nigra pars reticulata (SNr) to cortical stimulation. **A**. A sagittal section showing electrode traces with DiI. **B**. A map, raster plot, and PSTH of a typical SNr neuron with class 1 response (a triphasic response). **C**. A typical neuron with class 2 response (monophasic excitatory response). **D**. Z-scored PSTH of all responsive SNr neurons clustered by K-means clustering (top) and the mean PSTH of each cluster (bottom). Colors correspond to cluster numbers. The thickness indicates the numbers of neurons in the cluster. **E**,**F**. Examples of SNr neurons that were classified as class 1 but also received class-2-like monophasic excitatory input. **G**,**H**,**I**. Neurons showing a very fast excitatory monophasic response (**G**), monophasic inhibitory response (**H**), and inhibition-excitation pattern (**I**). **J**. Average input maps of neurons with class 1 (left) and class 2 (right) responses in SNr. **K**. An inhibitory response map of cluster #1 neurons in SNr. **L**. Time courses of the responsive areas. **M**. The same analysis as **D**, but separated according to recording location. Black and gray circles indicate triphasic and inhibition-excitation patterns, respectively. **Fig 4-1** is available for additional analysis on the SNr data.

**Figure 4-1.**
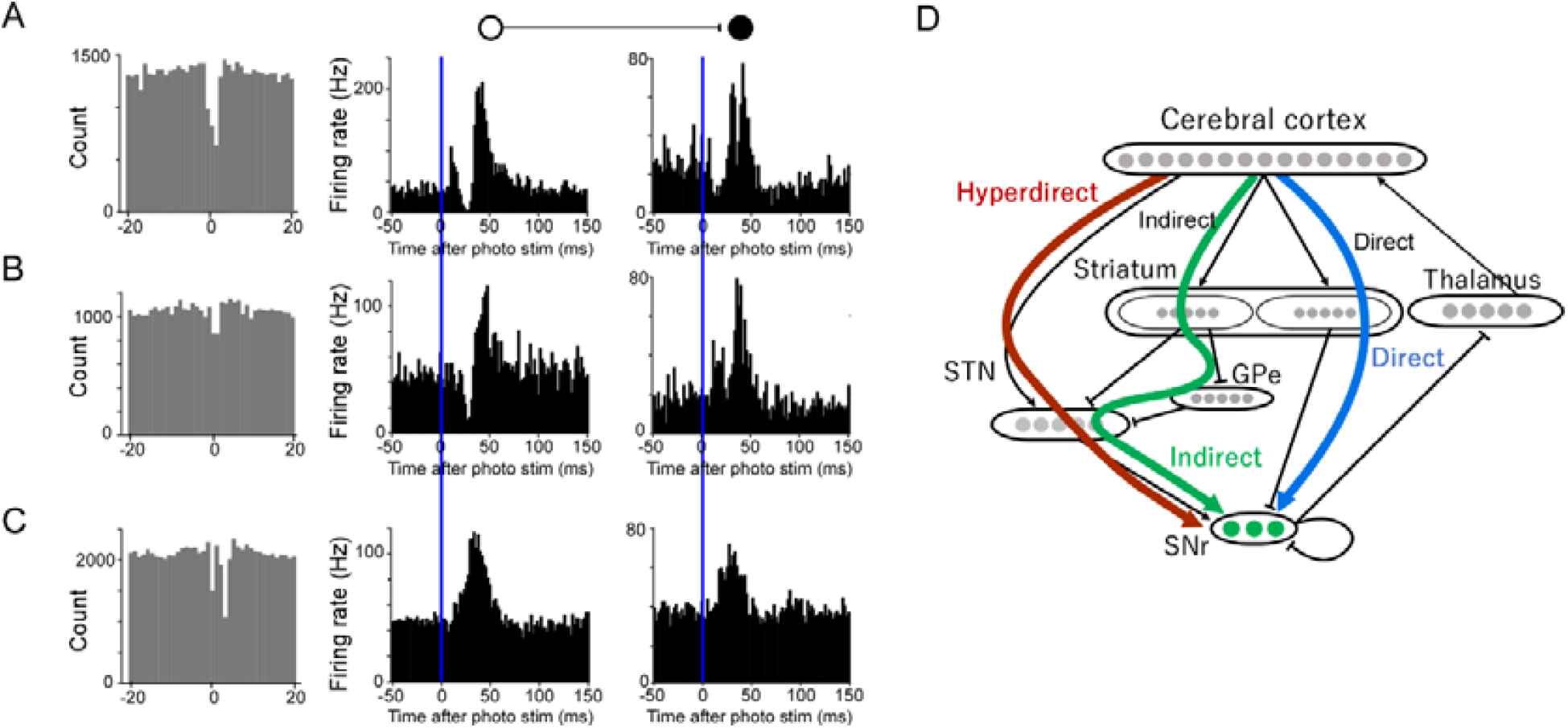
Hyperdirect, direct, and indirect pathways are explained by basal ganglia, not by synaptic interaction within the SNr. **A**. An example of cross-correlogram showing inhibitory interaction between simultaneously recorded pair of SNr neurons. The synaptic interaction does not explain the PSTHs. The PSTHs were obtained with the responsive pixels that shared by the two neurons. **B**,**C**. Examples of the other SNr neuron pairs. **D**. Schematic of the hyperdirect, direct, and indirect pathways.

We further investigated whether neurons with class 1 and class 2 responses are two completely different groups of neurons or whether a single neuron has both properties. One neuron with a class 1 response showed the triphasic response only when the frontal area was stimulated but showed a monophasic excitatory response when more posterior cortical areas were stimulated (**Fig. 4E**). Another neuron also showed a mixed response; however, class 1 and 2 responses were induced by the posterior and anterior part of the receptive field, respectively (**Fig. 4F**). These data are consistent with the idea that SNr can integrate information from different cortical areas with distinct patterns of activity transmission.

Some neurons only showed a fast excitatory response that had similar time courses to the first excitatory response of the class 1 triphasic response (**Fig. 4G**), an inhibitory response similar to the second inhibitory phase of the triphasic response (**Fig. 4H**), and an inhibitory-excitatory response pattern similar to the second and third phases of the triphasic response (**Fig. 4I**). Thus, the activities of neurons in SNr were characterized as based on the triphasic response, of which parts were missing (Nambu et al. 2023).

We further examined the dependencies of the response patterns on the locations of stimulation and recording. The excitatory source areas for neurons with class 1 and 2 responses distributed over a wide area of M2, M1, and S1 (**Fig. 4J**). Cluster #1 neurons in SNr also received inhibitory input from wide cortical areas including M2, M1, and S1 (**Fig. 4K)**. In clusters #1, #2, and #3, the sizes of the excitatory and inhibitory receptive fields were 3–4 mm^2^ and 1–2 mm^2^, respectively (**Fig. 4L**), supporting the center-surround model (Nambu et al. 2000). By contrast, more triphasic or inhibition-excitation responses were observed in the lateral SNr (>2 mm lateral from bregma) than in the medial SNr (p = 0.014, χ^2^-test; **Fig. 4M**). Thus, response patterns in SNr depend more on the location of the recording than the stimulus. Of note, the medial SNr may also receive more triphasic input from higher-order regions, such as the prefrontal cortex and orbitofrontal cortex (Maurice et al. 1999), which were not stimulated in our experiment.

We investigated what anatomical pathways each pathway is composed of. Cross-correlograms of SNr neurons showed rare inhibitory interactions (**Fig. 4-1A**). However, this inhibition does not explain the pattern of cortical responses to stimuli. The lateral inhibition inside the SNr does not seem to be large enough to shape the response patterns to cortical input. Based on previous studies (Fujimoto and Kita 1992; Nambu et al. 2000; Nambu, Tokuno, and Takada 2002; Sano et al. 2013), our data is consistent with the following circuit. The hyperdirect pathway runs from the cerebral cortex to the subthalamic nucleus to the SNr and is composed of fast transient excitation. The direct pathway inhibits SNr via the striatum. The indirect pathway activates the SNr after the hyperdirect and direct pathway inputs. The direct pathway can disinhibit the thalamus and cortex and makes a positive feedback loop between the cerebral cortex and basal ganglia. The hyperdirect pathway and indirect pathway localize the direct pathway spatiotemporally, resulting in a feedback circuit in which the cerebral cortex is continuously gated by the SNr (**Fig. 4-1B**).

### Retrograde tracing experiment

To determine which thalamic nucleus the SNr or CN neurons at the location we recorded project to, we injected AAV-based retrograde tracer (Tervo et al. 2016) (AAVretro-CAG-tdTomato) into the thalamus of Thy-1 ChR2 transgenic rats (**Fig. 5A**). Injections into the VL resulted in many tdTomato-labeled neurons in the bilateral CN but only a few in SNr (**Fig. 5C–F**). Within CN, labeled neurons were observed in the medial, interposed, and lateral CN bilaterally, but was brighter in the contralateral than the ipsilateral CN. In the contralateral cerebellum, axons collateral of CN neurons in the white matter and granule cell layer were also clearly labeled (**Fig. 5G.H**). We also confirmed tdTomato-positive neurons in layers 5 and 6 of the motor cortex (**Fig. 5I,J**). After injection of VM, tdTomato-labeled neurons were observed in the ipsilateral SNr at 2.1–2.6 mm lateral to the bregma (but not in the medial SNr) and in medial, interposed, and lateral CN bilaterally (**Fig. 5K–O**). After injection of VA, tdTomato-labeled neurons were observed in the ipsilateral SNr at 1.4–2.6 mm lateral to the bregma and in medial, interposed, and lateral CN bilaterally (**Fig. 5P–T**). In all cases, contralateral CN were more densely labeled than ipsilateral CN, whereas contralateral SNr was not labeled. These results indicate that the neural activity of the CN propagates to the VL, VA, and VM, and that the activity of the lateral SNr propagates to the VM and VA, which is largely consistent with previous studies (Kaneko 2013; Kuramoto et al. 2009; Gornati et al. 2018; Kuramoto et al. 2015).

**Figure 5.**
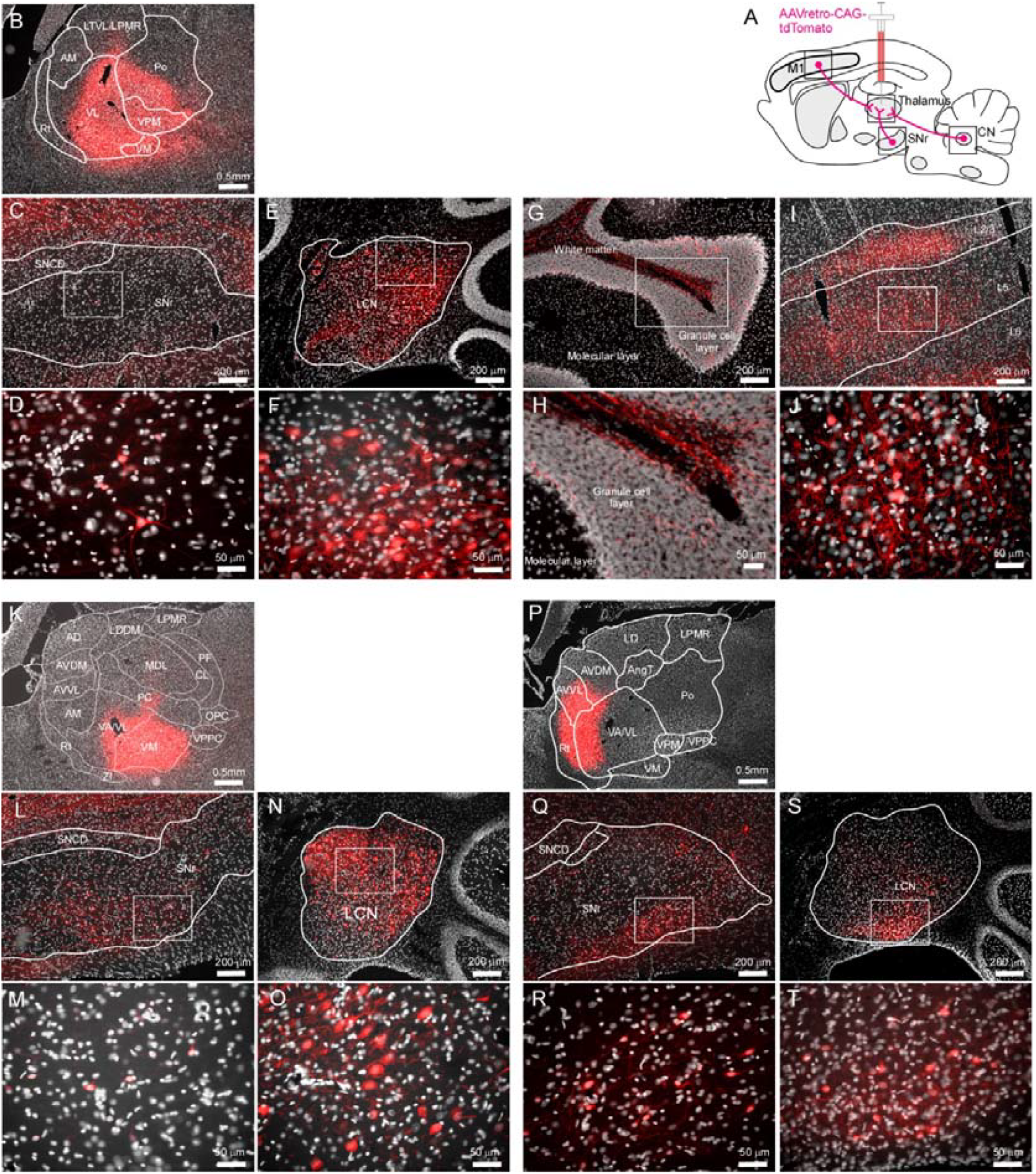
A retrograde tracer revealing thalamic projections from substantia nigra pars reticulata (SNr) and cerebellar nuclei (CN). **A**. Schematic of tracer experiments. **B**. Sagittal section 2 weeks after injection of AAVretro-CAG-tdTomato into the VL nucleus of thalamus. White is DAPI fluorescence showing localization of DNA. **C**. The tdTomato fluorescence around the ipsilateral SNr. **D**. Expanded image corresponding to the rectangle in **C. E**. The tdTomato fluorescence around the contralateral CN (LCN, lateral cerebellar nucleus). **F**. Expanded image corresponding to the rectangle in **E. G**. Axon collateral of CN neurons into the cerebral granule cell layer. **H**. Expanded image corresponding to the rectangle in **G. I**. The tdTomato fluorescence around the ipsilateral motor cortex. **J**. Expanded image corresponding to the rectangle in top panel. **K-O**. The same images as **B-F** of an injection experiment to the VM nucleus of thalamus. **P-T**. The same images as **B-F** of an injection experiment to the VA nucleus of thalamus.

### Cerebellar output to the thalamus, motor cortex, and striatum

The cerebellum and basal ganglia have direct connections that do not involve the cortex via thalamus (Bostan and Strick 2018; Yoshida et al. 2022; Ichinohe, Mori, and Shoumura 2000). The excitatory response of the CN when stimulated by the cerebral cortex might be transmitted to the striatum via the thalamus with a short latency (Chen et al. 2014). To investigate the fast propagation of activity from the cerebellum to the basal ganglia, we conducted electrophysiological recordings of the CN, motor cortex, centrolateral (CL) and ventrolateral (VL) thalamic nuclei, dorsolateral striatum (DLS), and dorsomedial striatum (DMS) during cerebellar stimulation (**Fig. 6**). We analyzed the neurons significantly increased the firing rate within 10 ms after photostimulation. Oblique laser incidence from behind the cerebellum through the transparent skull produced a very fast latency response (< 10 ms) in response to photostimulation (3.3 ± 0.40 ms, 8/37 neurons) (**Fig. 6G-I**). We confirmed that the CN neurons of the same transgenic rats express ChR2. The area where this response was detected covered the size of the CN (**Fig. 6G**), indicating direct stimulation of ChR2 on the soma or axons extending into the cerebellar cortex (**Fig. 5E**). We found evoked activity with a latency of < 10 ms in the VL (**Fig. 6J-L**; 5.0 ± 0.11 ms), CL (**Fig. 6M-O**; 5.4 ± 0.27 ms), motor cortex (**Fig. 6P-R**; 5.6 ± 0.27 ms) and striatum (**Fig. 6S-U**; 5.1 ± 0.40 ms; DLS or DMS). This indicates that the fast latency activity in DLS and DMS could be caused through the CN-thalamus-striatum pathway without going through the motor cortex. However, only 4.8% (6/126) of DLS and 4.1% (7/172) of DMS neurons responded to CN stimulation with a latency of < 10 ms, indicating that this CN-to-striatum activity propagation occurred only in a limited number of neurons. In fact, > 100 ms long-lasting activity was not often observed in SNr but in CN upon cerebral cortex stimulation. Thus, although this pathway has important functions in many aspects (Yoshida et al. 2022; Ichinohe, Mori, and Shoumura 2000; Alexander, DeLong, and Strick 1986), we will not consider it in the present study.

**Figure 6.**
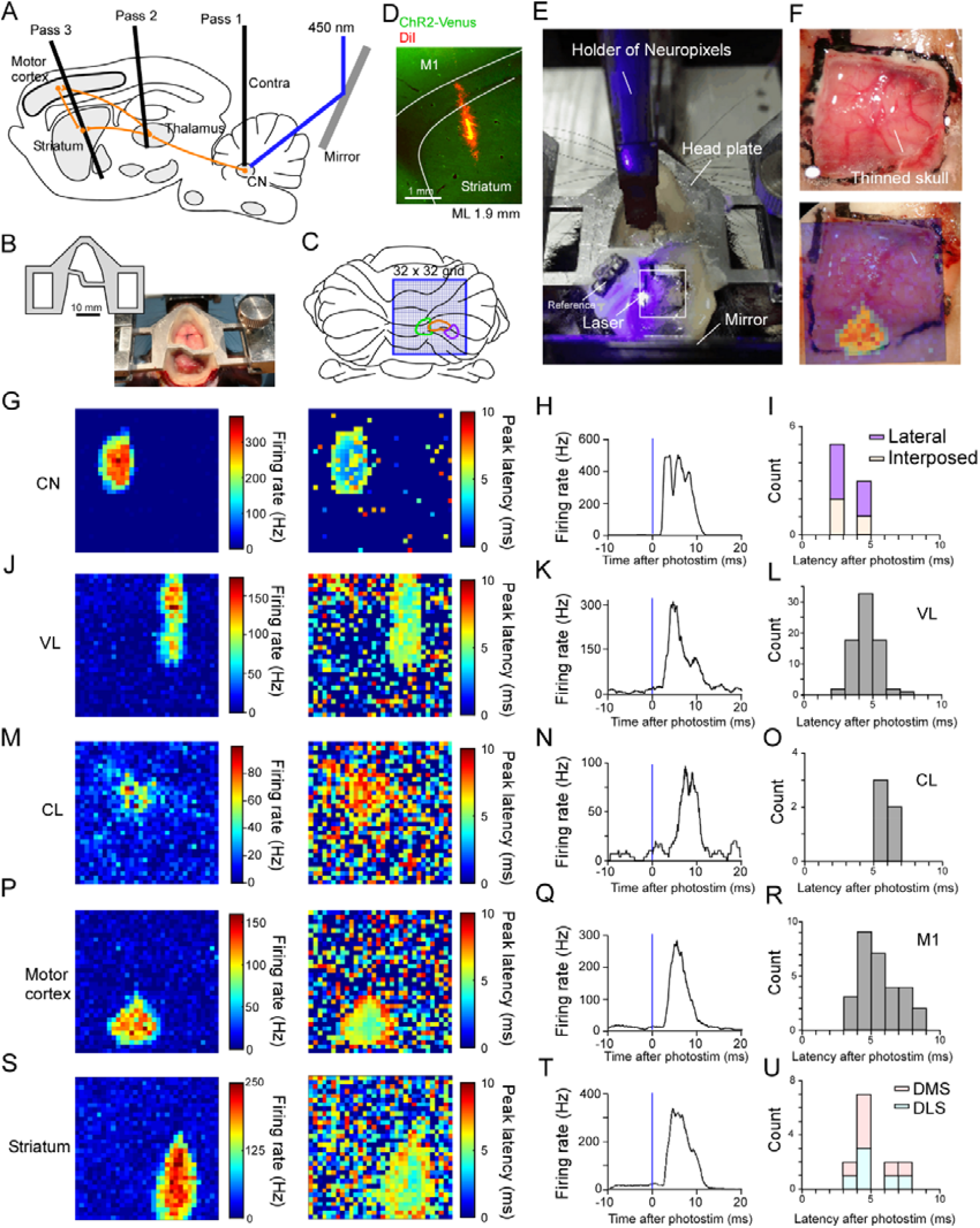
Activity propagation from the cerebellum to the striatum. **A**. Approximate positions of recording pass 1 for CN, pass 2 for the thalamus, and pass 3 for the motor cortex and striatum. The mirror was positioned behind the cerebellum to direct the laser to CN. **B**. A headplate optimized for this experiment. **C**. The cerebellum viewed from the angle of laser incidence. The blue frame indicates the photostimulation area. The approximate position of the lateral (purple), interposed (orange), and medial (green) nucleus were overlayed. **D**. Representative fluorescence image used to identify the location of the recording electrode. **E**. A photograph during photostimulation of the cerebellum and recording at the contralateral cerebral cortex. **F**. A photograph of transparent skull on the cerebellum (top). A response map was overlayed (bottom). **G**. Response map (left) and map of the peak latency (right) of a neuron recorded in interposed CN. **H**. The PSTHs of the receptive fields. **I**. Latency histograms of all neurons recorded in CNs. **J-L**. The same analysis as **G-I** for neurons in VL. **M-O**. The same analysis as **G-I** for neurons in CL. **P-R**. The same analysis as **G-I** for neurons in motor cortex. **S-U**. The same analysis as **G-I** for neurons in striatum.

### Hypothesis on brain-wide temporal integration

As CN→thalamus is excitatory and SNr→thalamus is inhibitory, increases in CN activity and decreases in SNr activity positively affect the thalamus. Notably, the excitation of class 2 response and the transient inhibition of the second phase of class 1 SNr response should coincidently activate the thalamus, as both of them two occur 20–30 ms after cortical stimulation (CN, **Fig 2D;** SNr, **Fig 4D**). Moreover, these two receptive fields almost overlapped (**Fig. 2I; Fig. 4J**). Therefore, a few tens of ms after the activity of a particular cortical area, signals that have passed through the SNr and CN circuits feedback synchronously to the cerebral cortex via the thalamus. We hypothesized that this synchronous positive feedback is the key to understand the brain-wide dynamics and learning. How the influence returns to the thalamus and cortex via the cerebellum and basal ganglia cannot be clarified experimentally because it is expected to be obscured by the mixture of various signals in the thalamus. Accordingly, in the next section, we performed computer simulations to investigate how this brain-wide synchronous circuit works in a coordinated manner.

### Spiking neural network reproducing representative activity of CN and SNr

Next, to investigate whether the activity propagation through these six pathways aligns with connectome data, we constructed spiking networks for the basal ganglia and cerebellum (**Fig. 7A,B; Fig. 7-1A; Table 2**). For the basal ganglia, we built a spiking neural network based on the connectome data using the Izhikevich model (Izhikevich 2003) (**Fig. 7-1B;** see **Methods**). This network contained the striatum, subthalamic nucleus, external globus pallidus, SNr, and thalamus (VL, VM, and parafascicular nucleus (PF)). The model reproduced a triphasic response to cerebral cortical stimulation in SNr neurons. To estimate which pathways from the cerebral cortex to SNr contribute to each phase of the triphasic response, we conducted virtual blocking experiments of synaptic connections (**Fig. 7C**). Blocking the hyperdirect pathway (i.e., monosynaptic input to the subthalamic nucleus from the cerebral cortex) eliminated the initial short excitatory response. Blocking the direct pathway abolished the second phase and weakened the third phase. Blocking the indirect pathway weakened the third phase.

**Figure 7.**
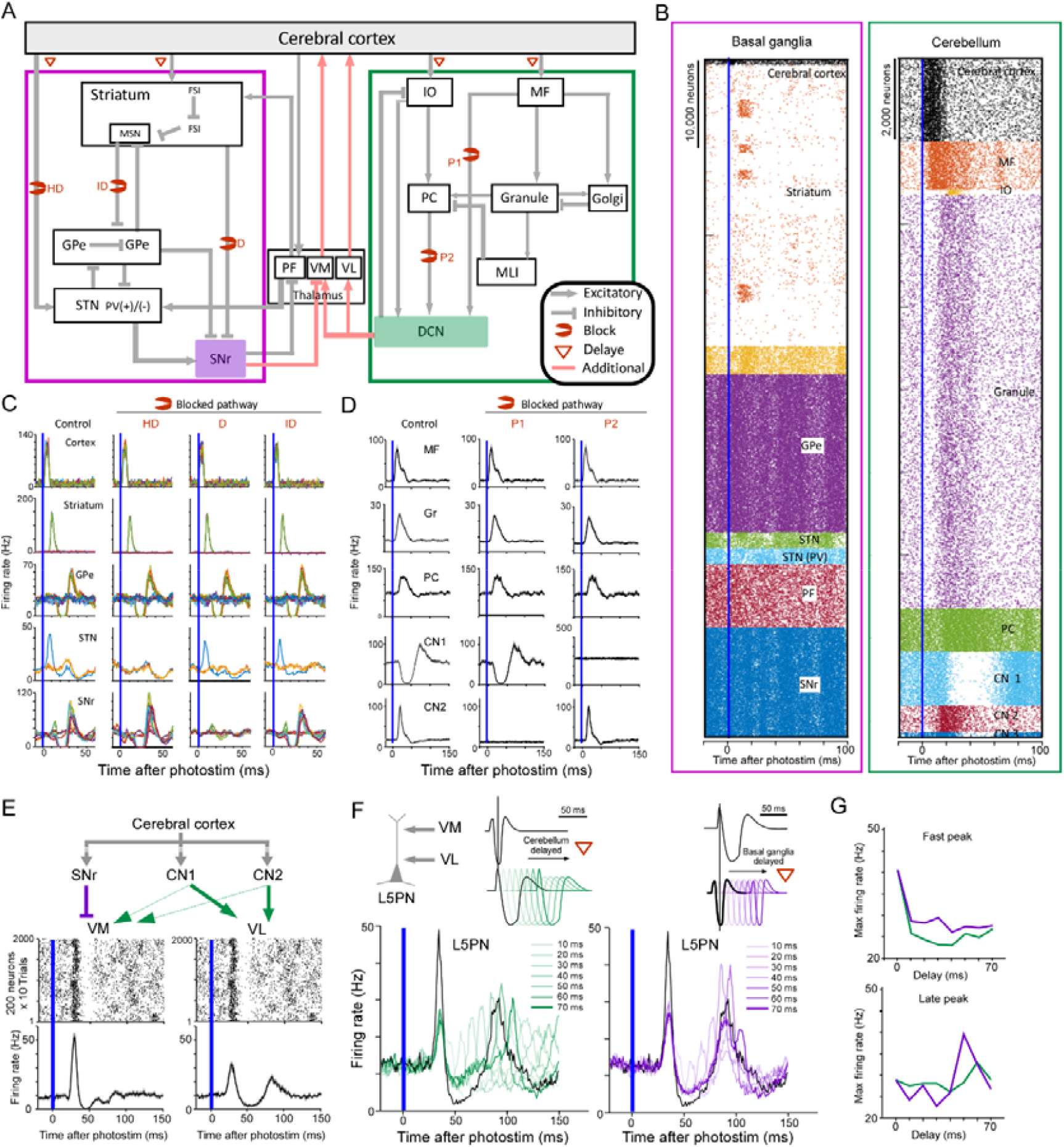
Simulation of spiking neurons based on the connectome data. **A**. Circuit schematic of the basal ganglia and cerebellum. In the basal ganglia, submodule-level details were constructed based on connectome data. Both share input from the cerebral cortex and output to the thalamus. **B**. Raster plot of simulated neurons before and after stimulation (single trial). **C**. PSTHs of major neurons in the basal ganglia under control conditions and after blocking the hyperdirect, direct, and indirect pathways. Colors indicate submodules. **D**. PSTHs of major neurons in the cerebellum under control conditions and after blocking MF-CN synapses (P1) and PC-CN synapses (P2). **E**. Simplified circuit for the simulation of VM and VL (top). Raster plots and PSTHs of VM and VL neurons (bottom). **F**. L5PN was assumed to receive both inputs from VM and VL (top left). The firing rate of L5PN was simulated when the signal from cerebellum was delayed (left), or when the signal from basal ganglia was delayed (right). **G**. Maximum firing rate of the L5PN at the fast peak (<50 ms) as a function of the delay-time of basal ganglia (purple) and cerebellum (green) (top). The same analysis for the late peak (>50 ms) (bottom). **Fig 7-1** is available for additional data and analysis.

**Figure 7-1.**
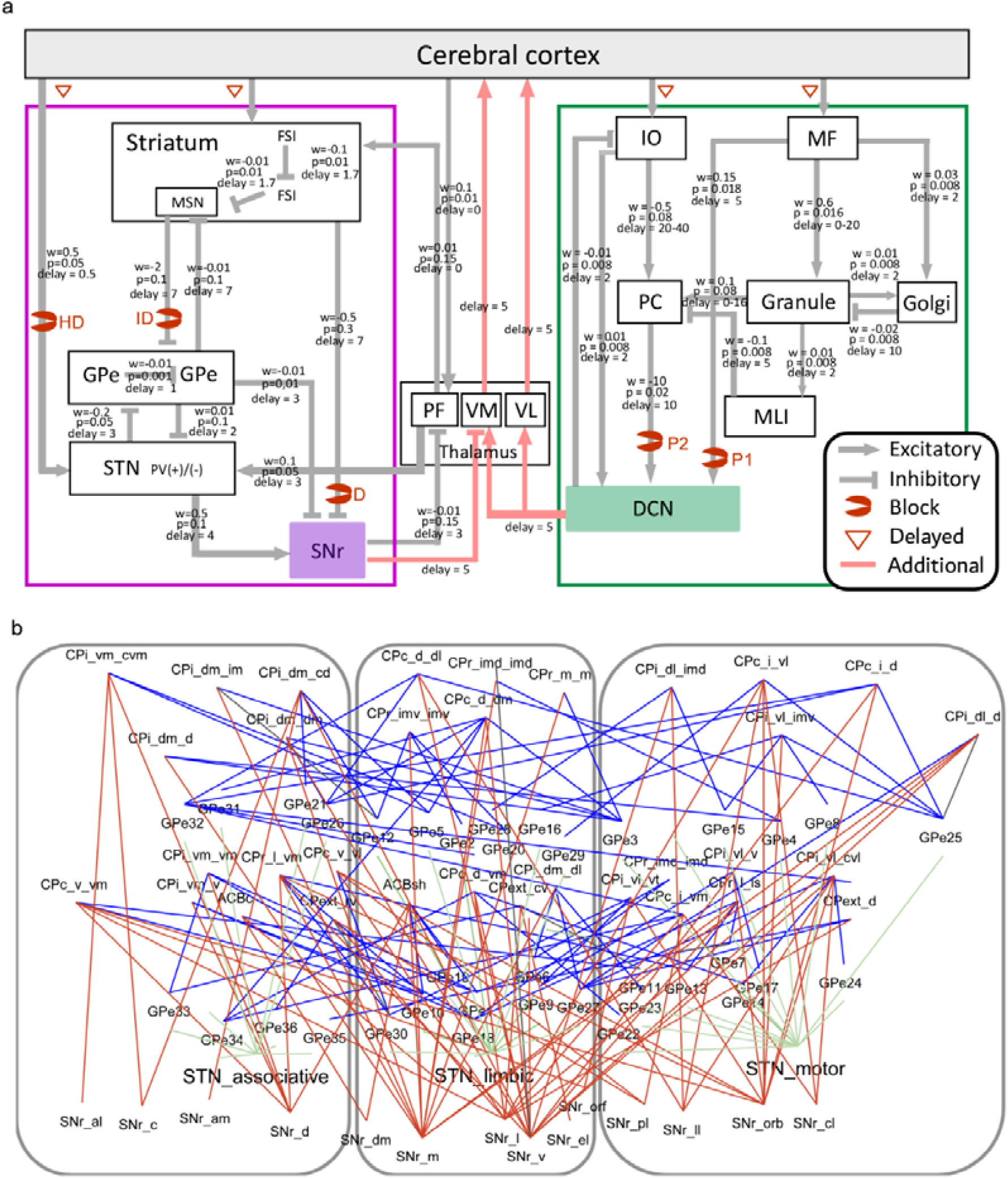
The network of cerebral cortex, cerebellum, and basal ganglia. **A**. A circuit diagram with parameters (w: weight in mV, p: probability, and delay in ms). **B**. Based on anatomical data, we constructed a detailed circuit consisting of 16 modules of striatum, 36 modules of external globus pallidus, 3 modules of subthalamic nucleus, and 14 modules of substantia nigra pars reticulata. Each line indicates a connection between subdivisions. The three rectangular frames indicate known clusters (Foster et al. 2021). Connections among CP (or ACB) (caudate putamen or nucleus accumbens, total 16 clusters), GPe (36 clusters), STN (3 clusters), SNr (14 clusters) were determined according to (Foster et al. 2021, Jeon et al. 2022, Hintiryan et al. 2016). Line colors indicate connection types.

Regarding the relationship between the cerebral cortex and cerebellum, it has been reported that the topography of the cortex is maintained particularly in the outputs from the pontine nuclei (Wu et al. 2023). However, this topography may be largely disrupted from granule cells to PCs (Pisano et al. 2021). Therefore, instead of creating a network model at a submodule level equivalent to that of the basal ganglia, we created a single module model for the cerebellum. We constructed a simulation consisting of MFs, granule cells, neurons in inferior olive, molecular layer interneurons, Golgi cells, PCs, and CN (**Fig. 7A,B; Table 2**). This model replicated the activity of class 1 and class 2 responses in CN (**Fig. 7D**). We conducted virtual blocking experiments as well. By blocking the projections from pontine nuclei to CN, only class 2 transient excitatory responses were eliminated. Blocking the synaptic functions from PCs to CN abolished class 1 responses. Therefore, CN neurons with class 1 inhibitory responses were inhibited by the PCs and then rebounded after inhibition in this model. Overall, our simulation replicated our electrophysiological results based on our current understanding of rodent brain circuits.

### Integration of two loop circuits during ongoing activity

To examine how the CN and SNr responses merged, we added projections from CN and SNr to the thalamus. Based on the results of tracer experiments, we assumed that VM receives both inputs from SNr and CN, while VL only received input from CN. We also assumed here that these thalamic neurons do not receive direct input from cerebral cortex. Approximately 30 ms after photostimulation, both thalamic activities were elevated (**Fig. 7E**). VL showed second peak of activity. To investigate how these thalamic activities can merge in the cerebral cortex, we simulated layer 5 pyramidal neuron (L5PN) receiving both inputs (Kaneko 2013; Hooks et al. 2013). Again, we assumed that this neuron does not receive input from surrounding cortical neurons. The L5PN showed triphasic responses, where the first excitatory signal was the summation of the direct pathway and CN phase 1 activities, the second inhibitory signal was the summation of the indirect pathway and CN phase 2 activities, and the third excitatory signal was CN phase 3. To investigate whether the synchronization of direct pathway and MF-CN pathway plays a role in information integration, we changed the axon conduction time of either corticostriatal and cortico-subthalamic projections or corticopontine and cortico-olivary projections (**Fig. 7F,G**). The exact synchronization at the timescale of 10 ms enhanced the summation of the CN and SNr outputs at the L5PN approximately 40 ms after the stimulation of cerebral cortex. Thus, our simulation reveals the importance of the temporal dynamics of outputs from the cerebellum and basal ganglia for information transmission back to the cerebral cortex via the thalamus. In the real brain, the interactions between thalamus and cortex that we have neglected would be complex. Synchronization of signals through the cerebellum and basal ganglia would be additive on top of such activity, thus controlling neurons in the cortex.

### Reservoir-based reinforcement learning model

Finally, we examined how the interaction of cortico-cerebellar and cortico-basal ganglia circuits contributes to learning. Recent studies have shown that in addition to dopamine neurons in the basal ganglia, complex spikes in cerebellar PCs reflect reward prediction error (Sendhilnathan et al. 2020; Kostadinov and Häusser 2022; Hoang et al. 2023; Heffley and Hull 2019; Thoma et al. 2008). This indicates that reinforcement learning should progress in both the basal ganglia and cerebellum. Thus, we investigated how reinforcement learning, which progresses in these different systems, merges coherently.

We built a model in which the activity of the cerebral cortex was represented by a reservoir of 800 units with a time constant of 20 ms and that produced complex dynamics in response to an auditory cue and reward inputs (**Fig. 8A**). For the basal ganglia, we assumed that the output is the result of convolving cerebral cortex activity with pathway-specific filters in the hyperdirect, direct, and indirect pathways (**Fig. 8B**, left). Similarly, for the cerebellum, the output was assumed to be the result of convolving the reservoir output with filters corresponding to CN phases 1, 2, and 3 (**Fig. 8B**, right). These six filters were made to tightly reproduce the experimental data.

**Figure 8.**
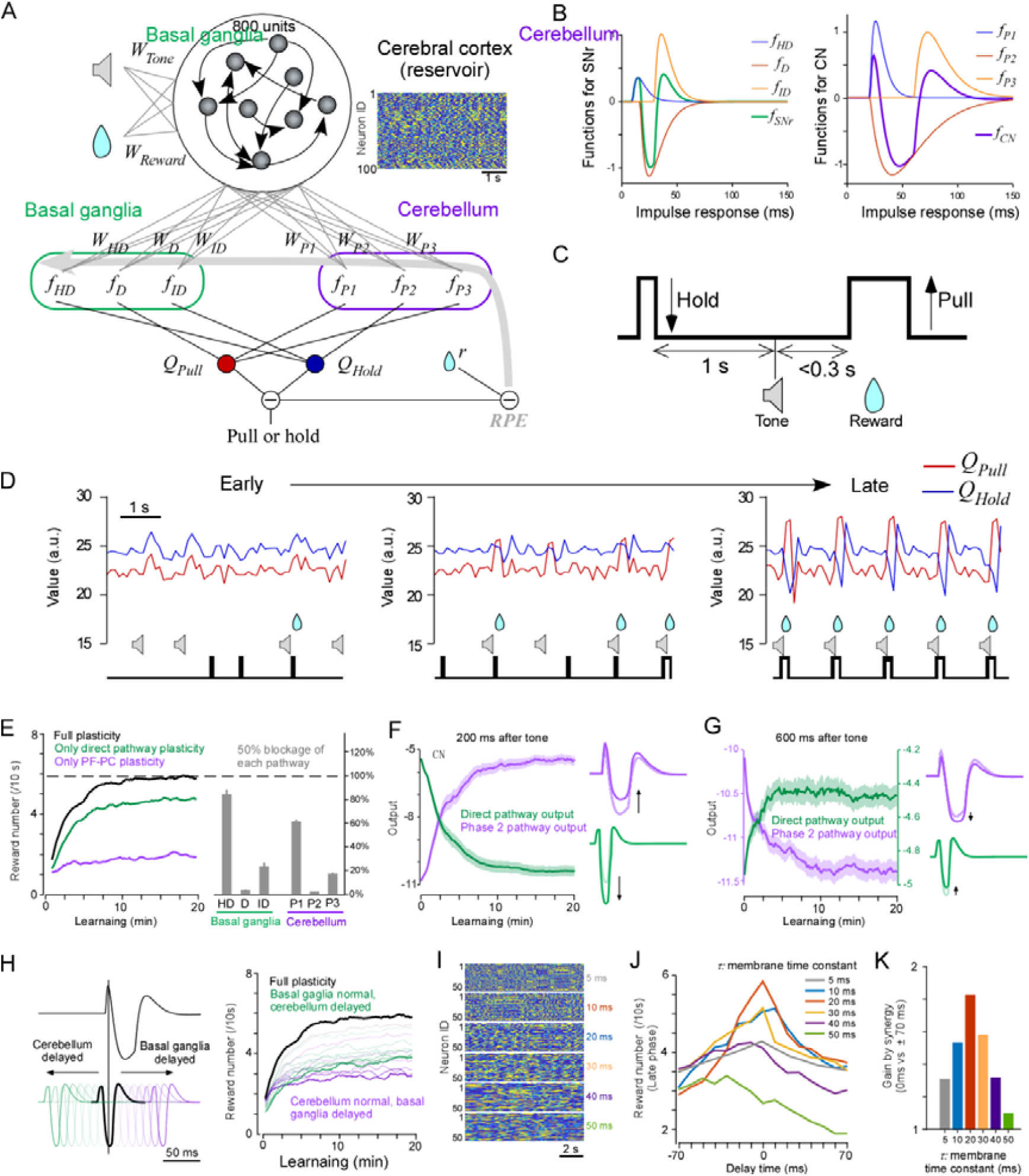
Reservoir-reinforced learning model of the cortex, cerebellum, and basal ganglia. **A**. The task structure. The agent is rewarded for holding the lever for 1 s followed by a sound cue and pulling the lever within 0.3 s. **B**. Double-exponential curves mimicking substantia nigra pars reticulata (SNr) (left) and cerebellar nuclei (CN) (right) activity. **C**. Schematic of the model. Cerebral cortex activity was simulated by a reservoir circuit that receives sound and reward inputs. The basal ganglia and cerebellum each integrate three synaptic strengths and three functions, each of which mimic the experimental data. The output from basal ganglia and cerebellum finally combined to a pull value and hold value. The difference between these values determined pull or hold once in 100 ms. Synaptic plasticity was induced by RPE. **D**. Examples of 6 s time series, pull or hold values, movement outputs, sound cues, and rewards in early to late learning stages. **E**. Learning curves under control conditions and after blocking synaptic plasticity in basal ganglia or cerebellum (left). The reward number in the late learning period after 50% blocking each pathway (right). HD; hyper direct, D; direct, ID; indirect, P1; phase 1, P2; phase2, P3; phase 3. **F**. Time course of direct pathway output (green) and P1 pathway output (purple) at 200 ms after tone presentation during learning. **G**. The same analysis as in f, but 600 ms after tone presentation. **H**. Schematic of signal delay(left). Changes in learning curves after delaying the conduction of corticostriatal or corticopontine projections. A 10 ms delay slowed the learning rate. **I**. Example of the activity of 50 units when the time constant tau of each unit in the reservoir was changed from 5 to 50 ms. **J**. Timescale of cortical neural activity indicated by colors affects the conduction-delay dependency of learning. **K**. Gain in learning rate when delay time was zero based on delay time plus or minus 70 ms. **Fig 8-1** is available for additional data and analysis.

**Figure 8-1.**
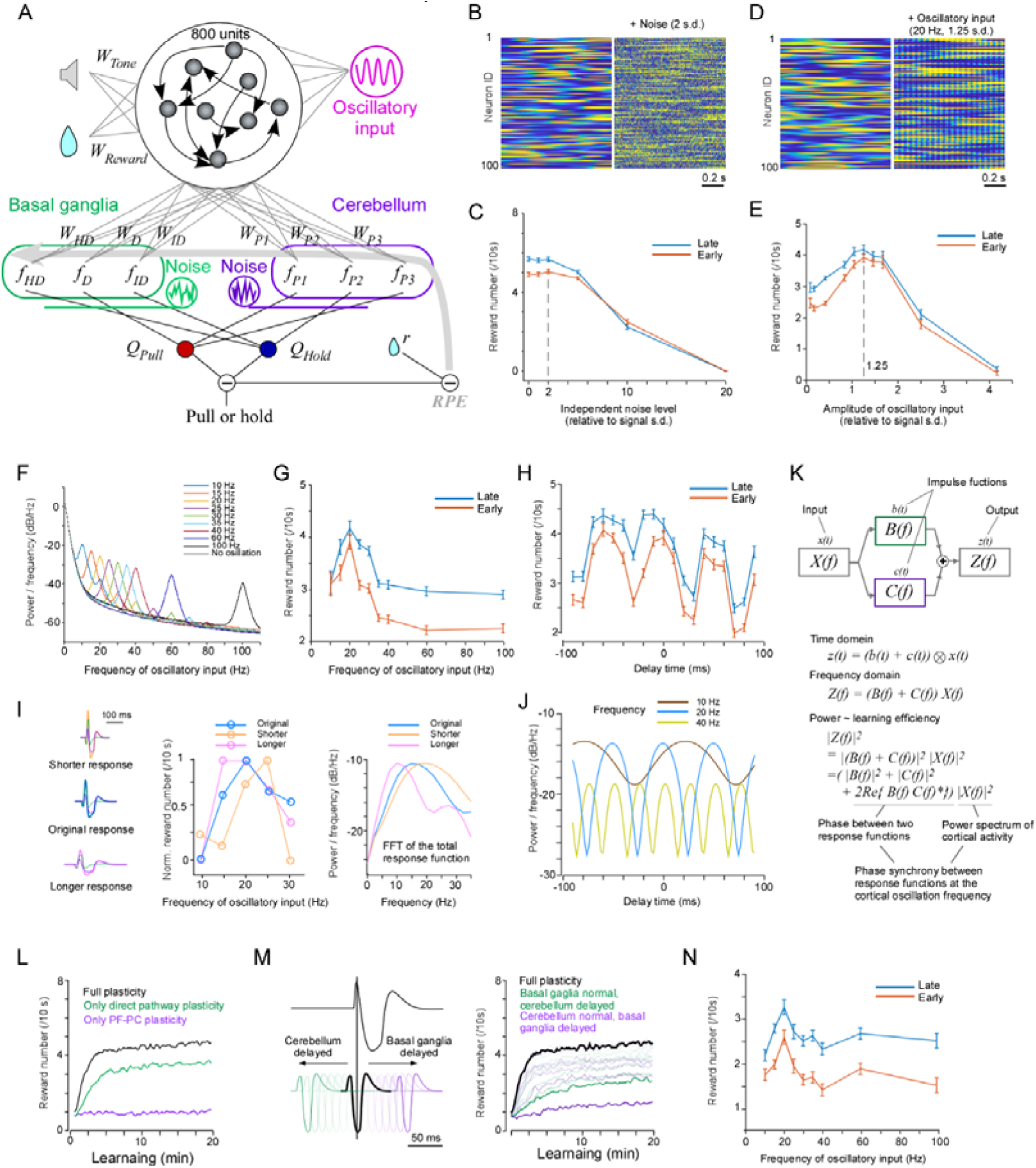
Noise and oscillatory input dependences of reinforcement learning. **A**. An illustration of independent noise input to the basal ganglia and cerebellum, or oscillatory input to the cerebral cortex (reservoir). **B**. Example of 100 neural activities of the reservoir circuit. Left, original neural activity. Right, neural activity when noise was added at twice the standard deviation of the neural activity. **C**. Number of reward acquisitions (10 s) during early and late learning periods when independent noise was applied to inputs to the basal ganglia and cerebellum. Learning efficiency did not deteriorate until the noise was about twice the standard deviation of the neural activity. **D**. Example of 100 neural activities of the reservoir circuit. **E**. Dependence of the number of reward acquisitions (10 sec) in the early and late learning periods on the amplitude of the beta oscillation. **F**. Power spectral density as a function of the frequency of the sine wave. **G**. Power spectral density of the number of reward acquisitions (10 sec) in the early and late learning periods. **H**. Time-delay dependence of the number of reward acquisitions (10 s) in the early and late learning periods with beta oscillations (20 Hz). A periodicity of about 50 ms, i.e., 20 Hz, was observed. This indicates that the phase difference between the basal ganglia and the cerebellum is involved in learning. **I**. ×0.8 (shorter, pink) or ×1.5 (longer, orange) time scale for CN and SNr response functions (**left**). Power spectrum density of the sum of CN and SNr responses (**middle**). Using original (blue), shorter (pink) or longer (orange) time scale filter, the number of rewards in the late learning period at each of the 10-30 Hz cortical oscillation frequencies was plotted against the cortical oscillation frequency (**right**). **J**. The power spectrum of the sum of all response functions when either the basal ganglia or cerebellar response function was delayed was plotted against the time delay of the basal ganglia or cerebellar response function. **K**. Parallel linear system consisting of the cortex (X), basal ganglia (B), and cerebellum (C). See main text for details. **L, M, and N**. The same analysis as in **Figure 8E, H** and **Figure 8-1G**, respectively, was performed by changing the time step for action from 100 ms to 50 ms. All showed qualitatively similar results.

Using this circuit, we trained a simple reinforcement learning task (**Fig. 8C**). The agent had to pull or hold a lever to get a reward. When it held the lever for 1 s, the tone cue was presented. The agent had to pull the lever within 0.3 s after the tone cue to get a reward. The decision to pull or hold the lever was made by comparing the pull value and hold value. The pull value was defined as the sum of the output results from the direct pathway, phase 1, and phase 3, since the activity of these pathways would enhance thalamic and cortical circuits leading to the pull behavior. The hold value was defined as the sum of the output results from the hyperdirect pathway, indirect pathway, and phase 2. These values were converted to lever pull probability using a sigmoid function, and based on this, the decision to pull or hold was made every 0.1 s. As pulling the lever requires force, a bias term was added to the sigmoid function to make pulling less likely. The final behavioral output, determined by integrating the outputs of the basal ganglia and cerebellum, was assumed to occur in the M1 via the thalamus. Synaptic plasticity occurred based on reward prediction error on corticostriatal synapses of the direct pathway and PF-PC synapses. Positive and negative reward prediction error (RPE) induced LTP and LTD in corticostriatal synapses, respectively, and LTD and LTP in PF-PC synapses, respectively (see **Methods**).

The simulation results show that the agent increased the reward acquisition rate during a 20 min training period (**Fig. 8D**). To investigate which components of the model are critical for successful learning, we first blocked the plasticity of either the basal ganglia or the cerebellum. This significantly reduced the rate of reward acquisition in both cases, suggesting that plasticity of both structures is important for learning in this model (**Fig. 8E left**). We then blocked 50% of the signaling in each pathway (**Fig. 8E right**). The blockage of any pathway inhibited learning, but in particular blocking either direct pathway in the basal ganglia or phase 2 pathway in the cerebellum abolished learning. Conversely, blocking the hyperdirect pathway in the basal ganglia and phase 1 pathway in the cerebellum had a relatively limited effect. These findings indicate that two pathways in which plasticity works play a particularly important role. Basal ganglia and cerebellar inputs contain independent components from other brain regions and periphery as well as cortical activity. We compared the number of rewards acquired in the early and late stages of learning by applying independent noise to the basal ganglia and cerebellar inputs. We found that learning was not impaired even when the standard deviation of the independent noise was twice the standard deviation of the signal from the cerebral cortex (**Fig. 8-1A-C**). This indicates the robustness of the cooperation between the basal ganglia and the cerebellum.

During the learning process, the pull value increased and hold value decreased immediately after the sound, whereas the pull value decreased and hold value increased during the holding period. In accordance with this, 200 ms after the tone presentation, the activities of the striatal direct pathway and cerebellar phase 2 pathway increased and decreased, respectively (**Fig. 8F**), whereas 600 ms after the tone presentation, these changes were milder (**Fig. 8G**). Thus, synaptic plasticity of both pathways occurred in a coherent manner to induce efficient behavior. Note that this model predicts that the triphasic activity patterns in response to cerebral cortex activity change from early to late learning stages (**Fig. 8F,G**).

To investigate whether the precise timing of the outputs from the basal ganglia and cerebellum contributes to learning, we delayed the output timing of the basal ganglia or cerebellum (**Fig. 8H**). Notably, delaying the output of the basal ganglia or cerebellum by only 10 ms reduced the reward acquisition rate. The greater the delay (up to 70 ms), the lower the reward acquisition rate. These findings suggest that the mutual timing of responses from the basal ganglia and cerebellum, as observed in experimental data, is optimized for cooperative reinforcement learning between the two structures. Thus, we investigated how this time-delay effect relates to the time-scale of cerebral cortex activity. We varied the time constants of neurons in 5 or 10 ms steps (from 5 ms to 50 ms) (**Fig. 8I**). When the time constant of a neuron was 10 ms, 20 ms, or 30 ms, delay-dependency was observed (**Fig. 8J,K**). These results suggest that delay-dependency requires a β- to γ-range timescale of cerebral cortex activity.

To investigate the relationship between cortical oscillation frequency and learning, we inputted oscillations as currents to the neurons, which constitutes the reservoir. When a 20 Hz sine wave was inputted, the global dynamics were similar to those of a sine wave with an amplitude 1.25 times larger than the original standard deviation (**Fig. 8-1D**). When the amplitude was varied, the number of rewards acquired in the early and late stages of learning was found to increase when oscillations of around 1.25 times the amplitude were given (**Fig. 8-1E**). Furthermore, when the oscillation frequency was changed while maintaining the amplitude of the oscillation, it was found that this enhancement effect occurred only with β-oscillations (20-30 Hz), not with γ oscillations (40-100 Hz) (**Fig. 8-1F, G**). Investigations of the time-delay effect showed that the number of reward acquisitions had 20-Hz oscillatory cycle, which indicate the successful learning depends on the coherence of cortical oscillation and sum of output from the basal ganglia and cerebellum (**Fig. 8-1H**). These results indicate that the facilitation of oscillatory frequency-dependent learning in the cortex is due to phase synchronization of feedback signals from the basal ganglia and cerebellum at the dominant frequencies. These results indicate that phase synchronization of cortical β-oscillations with those of the basal ganglia and cerebellum contributes to consistent reinforcement learning, which is consistent with previous studies on β-oscillations in the mammalian brain (Igarashi et al. 2013; Popa, Spolidoro, et al. 2013; Stein and Bar-Gad 2013).

To investigate in more detail what properties of the response functions of the basal ganglia and cerebellum are responsible for the oscillatory frequency dependence of the cortex, we performed simulations by varying the time scale of the response functions (**Fig. 8-1I**). When the response function was shorter in time (0.8 times) than the real data, i.e. when the feedback of cortical activity occurred on a shorter time scale, the frequency dependence shifted towards higher. Conversely, when the response function was longer in time (1.5-fold) and earlier than the real data, the frequency dependence shifted towards lower. Furthermore, this frequency dependence of learning matched well with the frequency dependence of the power spectrum of the sum of the six response functions at the respective time scales (**Fig. 8-1I, right**). This indicates that the oscillatory frequency-dependent facilitation of learning is contributed by the degree of agreement with the frequency of the feedback signals from the basal ganglia and the cerebellum. In addition, to investigate whether the results in **Fig. 8-1H** could also be explained by the power spectrum, we examined the power spectral density of those summations when the response functions of the cerebellum or basal ganglia were delayed. The delay time dependence of the power spectral density at 20 Hz was in sharp agreement with delay-dependent learning (**Fig. 8-1J**). In contrast, the power spectral densities at 10 Hz or 40 Hz were quite different. These results indicate that the facilitation of oscillatory frequency-dependent learning in the cortex is due to phase synchronization of feedback signals from the basal ganglia and cerebellum at the dominant frequencies.

Finally, to check whether the results of our reinforcement learning model were independent of the time step of the action, we changed the time step from 100 ms to 50 ms in our simulations. In line with this, we reduced the probability of pulling the lever in one decision by 1/2. This made the probability of pulling at 100 ms equal to the original probability. The results of blocking plasticity experiment, the time-delay dependence and the phase period dependence for the periodic input to the cortex were all qualitatively the same as for the 100 ms step (**Fig. 8-1L-N**). Thus, our reinforcement learning model is robust irrespective of time step.

## Discussion

In this study, we used optogenetics and electrophysiological recordings, which revealed that the response patterns of most single neurons in CN and SNr to cortical stimulation can be categorized into two classes. In CN, we observed inhibition followed by excitation (CN class 1) and transient excitation (CN class 2). In SNr, we observed a triphasic response (SNr class 1) and monophasic excitation (SNr class 2). The timescale of CN responses was slower than that of SNr responses. The effects on the thalamus in total were expected in the order of inhibition-excitation-inhibition in SNr and excitation-inhibition-excitation in CN. Notably, the MF-CN pathway of the cerebellum and the direct pathway of basal ganglia were synchronized. The response patterns could be reproduced by a spiking neural network model of basal ganglia and cerebellum based on currently available connectome data. We developed a model in which the cerebral cortex is the reservoir, and reward prediction error signal prompts LTP of striatal direct pathway neurons and LTD of parallel fiber synapses onto PCs. We found that cerebellar and basal ganglia dynamics aligned with experimental data with an accuracy of 10 ms facilitated cooperative reinforcement learning. Moreover, cortical β-oscillation was identified as critical for the successful learning. Overall, these data reveal a mechanism for the brain-wide synergy that underlies consistent reinforcement learning in the basal ganglia and cerebellum.

Previous computational studies have investigated the relationships among cerebral cortex, basal ganglia, and cerebellum (Houk and Wise 1995; Caligiore et al. 2019; Houk et al. 2007). However, our model with reinforcement learning is the first model based on realistic temporal response patterns of the cerebellum and basal ganglia in response to cerebral cortex activation. Our detailed analysis of temporal activity patterns revealed that β-range activity of the cerebral cortex enhances synergistic effects. Regarding oscillations, synergistic improvements were only observed for β-oscillations (20-30 Hz), not for γ-oscillations (40-100 Hz). The β-oscillation in the sensorimotor cortex has been observed during preparation and execution of voluntary movement (Stein and Bar-Gad 2013; Igarashi et al. 2013; Popa, Spolidoro, et al. 2013). Our model is also the first to assume that both the cerebellum and basal ganglia are engaged in reinforcement learning. Previously, it was assumed that the basal ganglia were responsible for reinforcement learning and the cerebellum for supervised learning, supposing a clear division of labor (Doya 2000, 1999). Our model proposes a novel mechanism of linking multiple plasticity events across distinct brain regions. That is, the synapses that are thought to most often undergo plastic changes in the basal ganglia and cerebellum are pathways that synchronously send back signals from the cerebral cortex. This parallel circuit can coincide by inducing plasticity on the direct pathway and PF-PC synapses, allowing reinforcement learning to proceed consistently.

Why does learning efficiency critically depend on cortical oscillation frequency and on the synergy of basal ganglia and cerebellar outputs at specific frequencies, even though our model simply assumes the summation of linear filters? Now, consider a parallel linear system with input and output time series *x*(*t)* and *z*(*t)* whose Fourier transforms are *x*(*f)* and *z*(*f)*, respectively, where *t* denotes time and *f* denotes frequency (**Fig. 8-1K**). The input and output correspond to cortical activity and decision probability, respectively. Let the response functions, i.e. impulse functions, of the basal ganglia and cerebellum be *b*(*t)* and *c*(*t)*, respectively, with corresponding Fourier transforms *B*(*f)* and *C*(*f)*. The composite response function is given by *b*(*f)* + *c*(*f)*, and by linearity, its Fourier transform is *B*(*f)* + *C*(*f)* The analysis regarding cortical oscillation suggested a close relationship between learning efficiency and the power spectrum of the output. This power spectrum |z(*f)*|^2^, which may be highly correlated with learning efficiency, can be expressed as

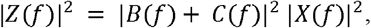

which expands to

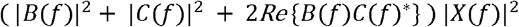

where *Re*{} denotes the real part and * indicates complex conjugate. Frequency shifts in response filters shown in **Fig. 8-1F,G,I** alter |*B*(*f)*|^2^ and |*C*(*f)*|^2^, potentially resulting in changes in learning efficiency. The cross term 2*Re*{*B*(*f*)C(*f*)*} depends on the phase difference between the response functions of basal ganglia and cerebellum, reaching a maximum when they are in phase and a minimum when they are in antiphase. This explains the time-delay dependency analyzed in **Fig. 8-1H,J**. Moreover, the phase synchrony at the frequency of cortical oscillation, which could enhance 2*Re*{*B*(*f*)C(*f*)*} | *X*(*f)*|^2^, is consistent with **Fig. 8-1I**. The close relationship between output power spectral density and learning efficiency may be due to the fact that a higher spectral density provides a broader dynamic range for synaptic plasticity to adjust the context-dependent modulation of behavioral outputs, thereby enabling robust learning. Therefore, during the learning process, to enhance the output power spectrum in the β band, the outputs of the basal ganglia and cerebellum may tend to become phase-locked at β-frequency range.

It is known that axons from VL and VM are distributed across the different layers of the cerebral cortex (Kaneko 2013). VM especially projects to apical dendrites in layer 1, whereas VL projects to the basal dendrites of L5 pyramidal neurons (Kuramoto et al. 2015). Synchronous inputs to tuft and proximal dendrites within 20 ms enhance nonlinear integration, leading to dendritic spikes and bursting (Ledergerber and Larkum 2012; Poirazi and Papoutsi 2020). In the motor cortex, cortico-spinal neurons may detect this coincidence and directly convert it into motor output. Thus, if cerebellar and basal ganglia outputs are synchronized with a precision of 20 ms, activity of VL and VM could facilitate strong motor output via dendritic spikes in the motor cortex. This necessary timescale is consistent with the timescale that we have identified to promote thalamic activity in simulation and coordinated reinforcement learning in our model (i.e. β-range). Dendritic spiking may also induce synaptic plasticity, and simultaneous active input from the cerebellum (Popa, Velayudhan, et al. 2013) and basal ganglia (Ueki et al. 2006) may facilitate circuit formation in the cerebral cortex. β band oscillation is found in the motor cortex, basal ganglia, and cerebellum (Courtemanche, Robinson, and Aponte 2013; Halje et al. 2019; Spencer and Ivry 2021; Middleton et al. 2008; Khanna and Carmena 2015; Morén et al. 2019; Igarashi et al. 2013; Leventhal et al. 2012). Taken together, we propose that the cerebellum, basal ganglia, and cortex acquire plasticity in a coordinated manner through β rhythm, resulting in consistent behavioral decisions and reinforcement learning throughout the brain.

Whether the mechanisms of consistent reinforcement learning that we modeled here explain the actual brain’s learning principle remains to be tested. As shown in **Fig. 8F**, excitatory transmission from CN to the thalamus and inhibitory transmission from the SNr to the thalamus will be larger and smaller, respectively, upon execution of the reinforced behavior after learning. By contrast, these transmissions will be smaller and larger, respectively, if the behavior does not need to be executed (**Fig. 8G**). It should be noted that CN and SNr activity in response to non-specific activation of the cerebral cortex will not change during learning. It is necessary to artificially stimulate cortical neurons to induce or suppress the behavior to be reinforced to verify these predictions. Moreover, optogenetic inhibition of PCs (whose activity is suppressed during the stimulation of cortical neurons that are involved in reinforced behavior) and optogenetic activation of striatal neurons in the direct pathway (whose activity is increased during the stimulation of cortical neurons that are involved in reinforced behavior) would cause the execution of the reinforced behavior via the thalamus and cerebral cortex. These experiments could be possible by combining an activity-dependent genetic switch (Wang et al. 2017; Lee et al. 2017), such as Cal-Light or FLARE (Lee et al. 2017; Wang et al. 2017) with tet-dependent. In addition, measuring the axonal activities from SNr and CN onto the thalamus during specific behavior would be useful. This could be done, for example, by expressing voltage sensor proteins with different fluorescence wavelengths in the SNr and CN and observing the thalamus through the GRIN lens. These axonal activities are predicted to be synchronized and phase-reversed in beta oscillations during the preparation or execution of a specific behavior.

We have found that CN responses are categorized into distinct class 1 and 2 activities. The excitatory inputs in class 2 were thought to be via MFs (Tsukahara, Korn, and Stone 1968; Hesslow, Svensson, and Ivarsson 1999; Wu, Sugihara, and Shinoda 1999); however, involvement via CFs (Sugihara, Wu, and Shinoda 1996) or disinhibition of PCs cannot be ruled out. Although many class 1 neurons did not show fast excitation, they can change to class 2 because MF-CN synapse potentiation can be induced by pairing with post-inhibitory rebound of CN neurons (Pugh and Raman 2008). As the MF-CN pathway is disynaptic, and the MF-granule cell-PC-CN pathway contains four synapses, the MF-CN pathway is considered faster. However, as indicated in **Fig. 2F** (* in the right panel), inhibition may precede excitation. Thus, even with signals starting simultaneously from the cerebral cortex, PCs may be able to alter the impact of the MF on the CN. CN phase 3 could represent post-inhibitory rebound excitation (Hoebeek et al. 2010; Aizenman and Linden 1999), excitation by internal circuits in CN, and reverberation by circuits not recorded in this study. A marked difference in the proportions of class 1 and class 2 in interposed and lateral CN could be due to the relative differences in MF and PC input. However, this requires further investigations in future studies.

Finally, we list below the limitations of this study and future directions. Convolution of the impulse response assumes a linear model; however, the brain is clearly not linear, and nonlinearities must be considered in many cases. We assessed the cerebellum and basal ganglia; nevertheless, other areas, such as the superior colliculus and sensory cortex, are also involved in decision making. We also disregarded the corticothalamic projection, even though the mechanisms by which inputs from the cerebral cortex merge with those from SNr and CN in the thalamus are most likely important. It is possible that rebound excitation of CN class 1 contributes to working memory maintained by the cerebro-cerebellar loop (Gao et al. 2018), which should be examined in the future. The mechanism by which the RPE is shared between the basal ganglia and cerebellum should also be clarified. As cooperation between the basal ganglia and cerebellum is important for facilitating learning throughout the brain, dysfunctions in one area may affect both areas. For example, diseases related to the basal ganglia, such as Parkinson’s disease, obsessive-compulsive disorder, and Tourette’s syndrome, may also be associated with the cerebellum as previously discussed (Bostan and Strick 2018; Caligiore et al. 2017; Li, Le, and Jankovic 2023). Thus, our findings may call for re-interpretation of various brain pathologies.

## Acknowledgements

We thank T. Shimada, R. Mizuno, Reiko Hira for animal husbandry and genotyping, M. Kawabata, A. Rios, T. K. Fujita, Sakairi, S.L. Smith, J.N. Stirman, S. Aoki, A. Funamizu K. Ishizu, S. Tsutsumi, M. Morishima, T. Ishikawa, T. Yamazaki, H. Mori for technical advices and discussion. We also thank the editor and reviewers for helpful comments. This work was supported by JP22wm0525007 (RH), JP24wm0625405 (RH), JP19dm0207089 (YI) from AMED, JP24H02156 (RH), JP22H02731 (RH), JP20K22678 (RH), JP21B304 (RH), JP21H05134 (RH), JP21H05135 (RH), JP16H06276 (YI), JP21H0524 2(YI), JP19H03342 (YI), JP23H02589 (YI), and JP20H05053 (YI) from MEXT/JSPS, JPMJCR1751 (YI) from JST, Nakatani Foundation (RH), Shimadzu Foundation (RH), Takeda Science Foundation (RH), Takeda Science Foundation (YI), The Precise Measurement Technology Promotion Foundation (RH), Tateishi Science and Technology Foundation (RH), and Research Foundation for OptoScience and Technology (RH).

## Code and data accessibility

The code of the spiking neural network, code of reservoir model, CAD files of the headplate, parts list, Zemax files for the scan-optogenetics system will be available upon publication. Electrophysiological data is available upon reasonable request.

