## Supplementary figures and images for "Synergistic reinforcement learning by cooperation of the cerebellum and basal ganglia" (from another version of this preprint: version 3, 2025-03-20)

### Extended Data 1

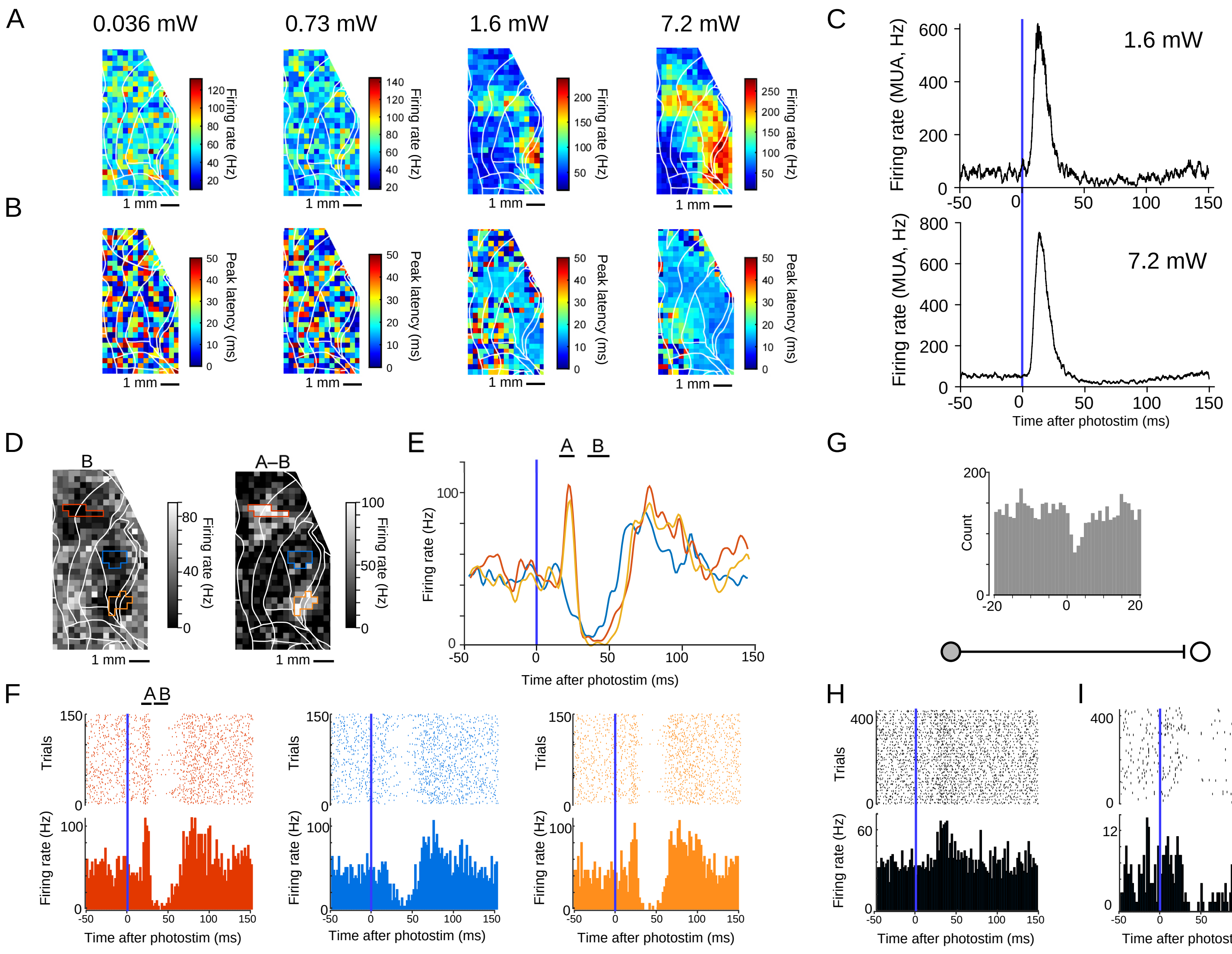

### Extended Data 2

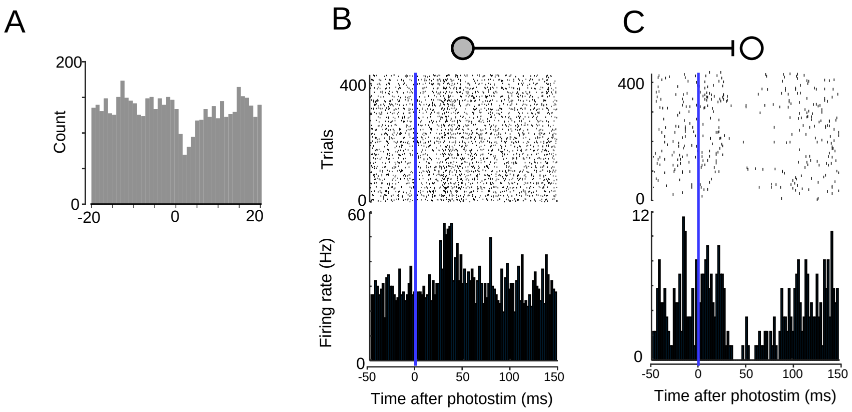

### Extended Data 3

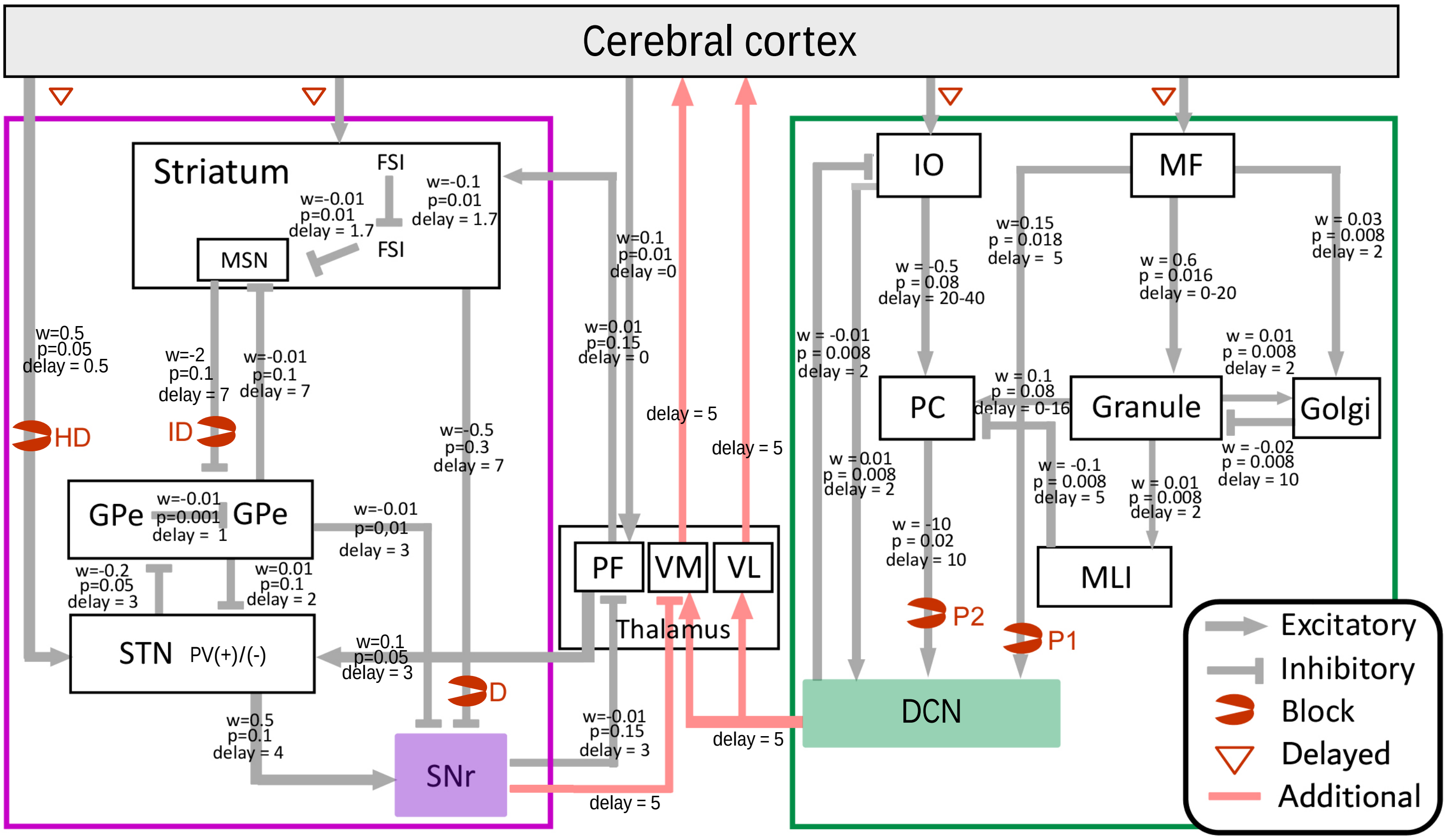

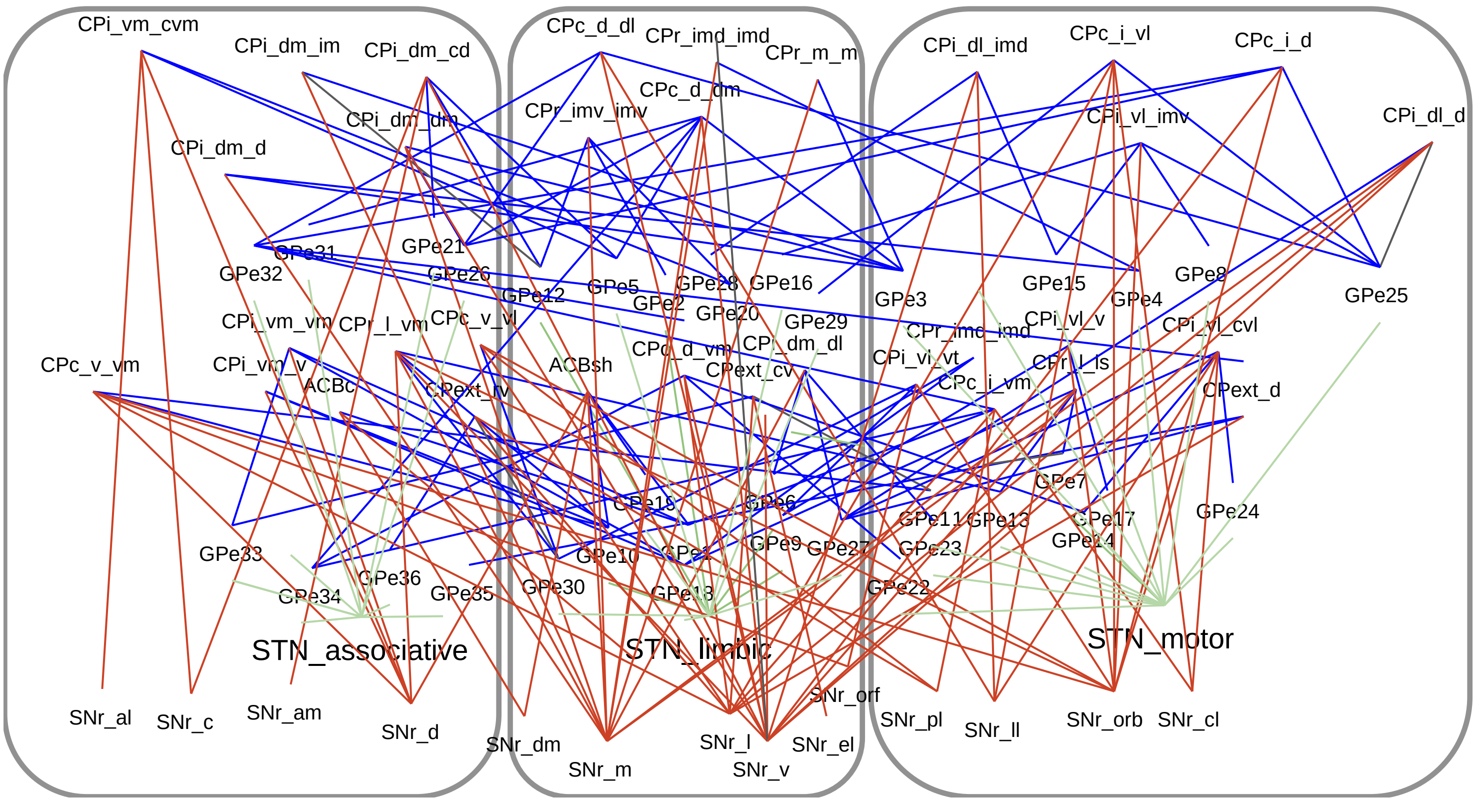

### Extended Data 4

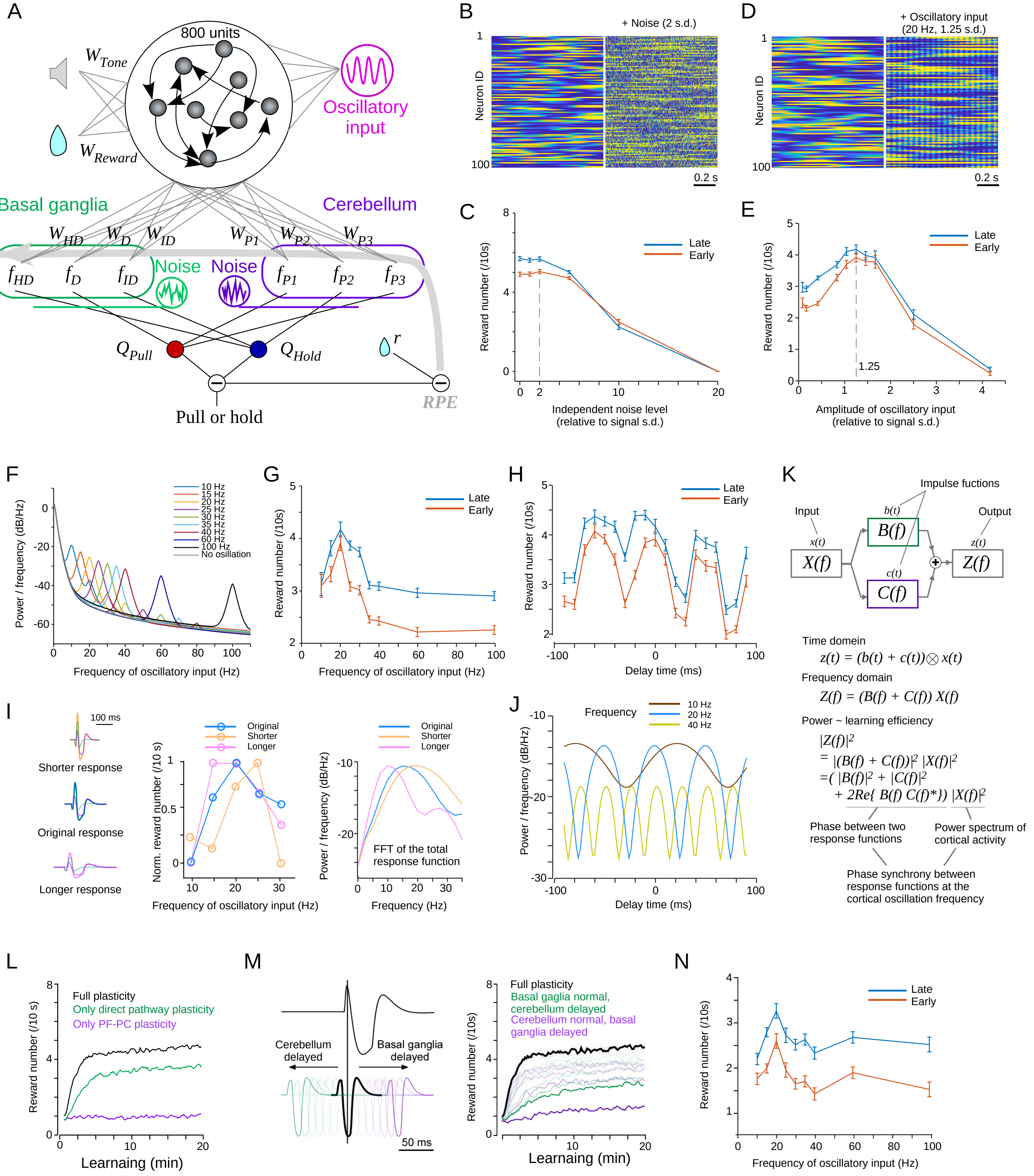
